# Inhibition of Lactate Dehydrogenase A (LDH-A) by Diclofenac Sodium Induces Apoptosis in HeLa Cells by Activation of AMPK

**DOI:** 10.1101/2023.10.02.560620

**Authors:** Avirup Malla, Suvroma Gupta, Runa Sur

## Abstract

Cancer cells exhibit a unique metabolic preference for choosing the glycolytic pathway over oxidative phosphorylation for maintaining the tumor microenvironment. Lactate dehydrogenase-A (LDH-A) is a key enzyme that facilitates glycolysis by converting pyruvate to lactate and has been shown to be upregulated in multiple cancers due to the hypoxic tumor microenvironment. Diclofenac (DCF), a non-steroidal anti-inflammatory drug, has been shown to exhibit anti-cancer effects by interfering with the glucose metabolism pathway. However, the specific targets remain unknown. Using *in-silico*, biochemical, and biophysical studies, we show that DCF binds to LDH-A adjacent to the substrate binding site and dose-dependently inhibits its activity in an allosteric manner in HeLa cells. Thus, DCF inhibits the hypoxic microenvironment and induces apoptosis-mediated cell death. DCF fails to induce cytotoxicity in LDH-A knocked-down HeLa cells, confirming that DCF renders its anti-mitotic effects via LDH-A inhibition. DCF-induced LDH-A inhibition alters pyruvate, lactate, NAD^+^, and ATP production in cells, and this could be a possible mechanism by which DCF inhibits glucose uptake in cancer cells. DCF-induced ATP deprivation leads to mitochondria-mediated oxidative stress, which results in DNA damage, lipid peroxidation, and apoptosis-mediated cell death. Reduction in intracellular ATP levels additionally activates AMPK, a sensor kinase, which further downregulates p-S6K, leading to apoptosis-mediated cell death. We find that in LDH-A knocked-down cells, intracellular ATP levels were depleted, resulting in the inhibition of p-S6K, implying the involvement of DCF-induced LDH-A inhibition in the activation of the AMPK/S6K signaling pathway.

## Introduction

Cancer cells remodel their metabolism for instant ATP production required for growth, survival, proliferation, and prolonged maintenance [1, 2]. Thus, cancer cells show a unique metabolic preference for choosing the glycolytic pathway rather than oxidative phosphorylation (OXPHOS) even in the presence of functional mitochondria and sufficient oxygen concentrations, which is known as the Warburg effect [1, 3]. This metabolic switch enables cancer cells to produce instant ATP and release lactic acid, resulting in the acidification of the tumor microenvironment. This makes them resistant to acid-induced cell toxicity, thus promoting unconstrained proliferation and invasion [4]. Another mechanism by which aerobic glycolysis could confer a proliferative advantage to cancer cells lies in the incomplete utilization of glucose. This allows the unused intermediates to be repositioned towards macromolecular biosynthesis, thus providing cancer cells with an abundance of carbon building blocks [5].

Lactate dehydrogenase (LDH) is a well-established enzyme (EC 1.1.1.27) that is mainly responsible for the reversible conversion of pyruvate to lactate using NADH as a cofactor [6, 7]. LDH is composed of four subunits and has two types of chains, namely A and B. LDH has five isoforms, namely A4, A3B1, A2B2, AB3, and B4, and all of them are either responsible for the forward reaction or the backward reaction [6, 7]. Lactate dehydrogenase-A (LDH-A, LDH-5, M-LDH-A) is mainly localized in muscle cells [8]. In the hypoxic microenvironment of tumor cells, the transcription factors HIF-1α and c-myc are upregulated, which further upregulates LDH-A which has been identified as one of the key enzymes in causing the Warburg effect [9, 10]. It has already been well established that reduction of LDH-A expression or inhibition of its enzymatic activity leads to inhibition of tumor initiation and cell proliferation [11]. Thus, LDH-A is considered a novel target for a variety of anticancer drugs [12, 13].

AMP-activated protein kinase, commonly known as AMPK, is a serine-threonine protein kinase that acts as a highly sensitive energy sensor that can sense intracellular energy fluctuations [14]. AMPK is activated by the depletion of intracellular ATP levels (more precisely by the increase in the ratio of AMP/ATP as well as ADP/ATP levels), glucose deprivation, hypoxia, and ischemic conditions [15]. Activation of AMPK inhibits the phosphorylation of its downstream target, S6K [16]. There is a very strong correlation between the activation of the AMPK signaling pathway and the induction of apoptosis in cancer cells [17, 18, 19]. AMPK activation also has a strong impact on cell growth and proliferation, survival, and protein synthesis [19, 20]. Many novel anticancer agents exhibit their efficacy by activating the AMPK signaling pathway in many cancer cell lines [18, 19, 21].

Diclofenac (DCF) is widely used as a painkiller due to its ability to decrease COX-2 expression in the prostaglandin production pathway [22)]. Aside from its anti-inflammatory properties, some data suggests that DCF has anti-mitotic properties in several cancer cell lines [23]. DCF seems to have antimitotic effects by interfering with the glucose absorption route, although the molecular mechanism underlying this action is unclear [24, 25]. This prompted us to identify the molecular target of DCF in the glucose absorption and metabolism pathway so that it may be repurposed as a targeted anticancer drug. Our results, for the first time, establish that DCF inhibits LDH-A in cancer cells and prevents its downstream effects such as glucose uptake, extracellular acidifications, the NAD^+^/NADH ratio, and ATP production. This lowering of intracellular ATP levels by DCF activates the AMPK signaling pathway, which is responsible for the initiation of apoptosis. Thus, this work identifies DCF as a lead anticancer molecule.

## Results

### DCF inhibits proliferation in HeLa cells

To find out the cytotoxic effect of DCF on cancer cells, HeLa cells (a human cervical cancer cell line) were treated with different concentrations of DCF, and a trypan blue exclusion assay was performed. Results showed that DCF treatment dose-dependently reduced cell viability in HeLa cells with an IC_50_ dose of 175 μM± 4.86 μM on 24 h of incubation (Fig. 1A). DCF time-dependently inhibited cell viability in HeLa cells when cells were treated with 100 μM DCF for different time points (Fig. 1B). Additionally, DCF treatment inhibited cell viability in other cancer cells, such as HCT-116 (a colon cancer cell line) (Fig. S1) and MCF-7 (a breast cancer cell line) (Fig. S2), suggesting that the results are not specific to cervical cancer cells alone but could be generalized to other cancer cells.

**Fig. 1.**
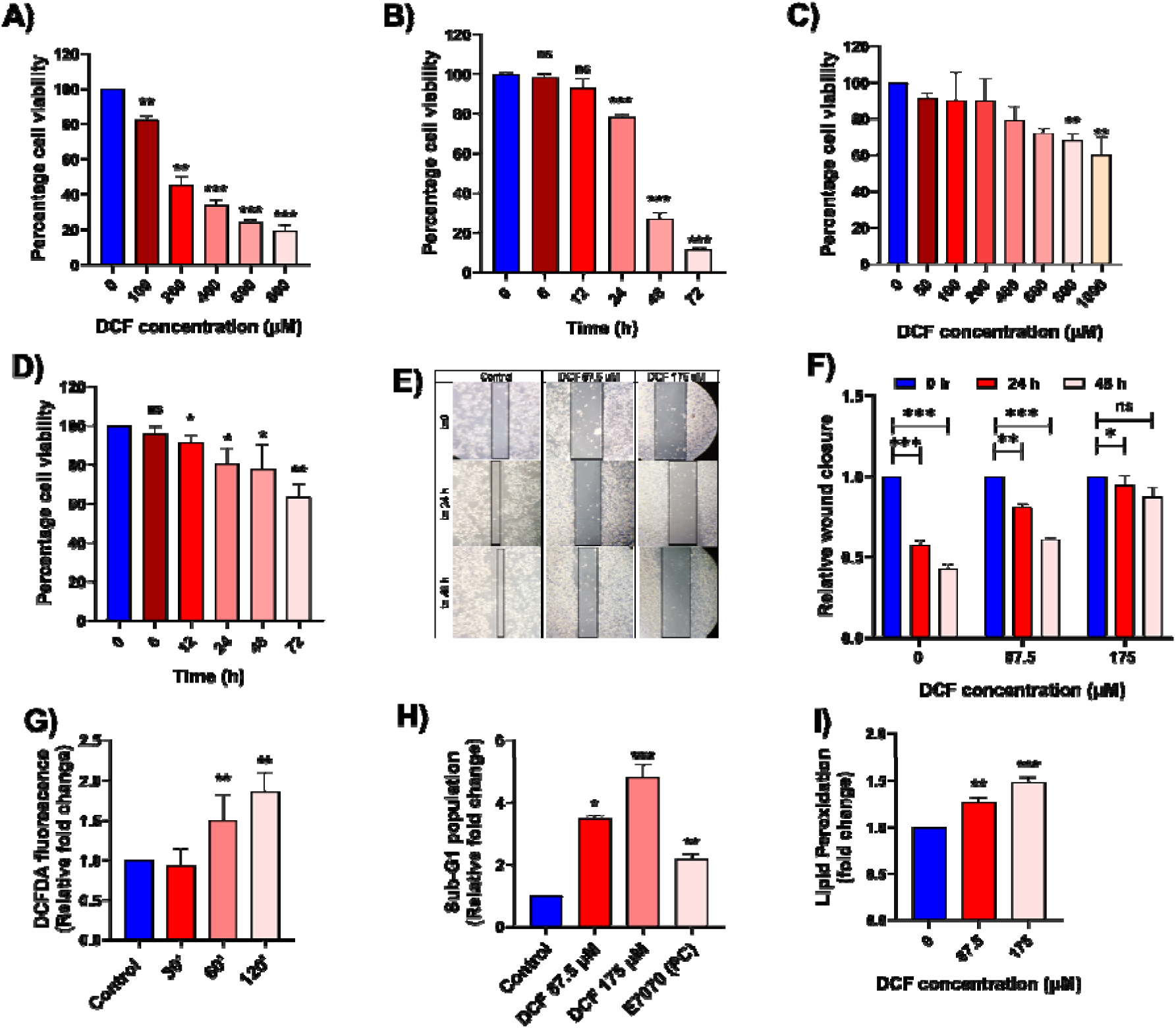
DCF inhibits cell proliferation in HeLa cells. (A) The percentage of cell viability was estimated using a trypan blue exclusion assay with different concentrations of DCF (0-800 μM) in HeLa cells for 24 h. (B) Cell viability was estimated with 100 μM DCF for different time points (0–72 h) in HeLa cells using the trypan blue exclusion assay. (C) Percentage cell viability was estimated with different concentrations of DCF (0–1 mM) in NKE cells. (D) Cell viability was estimated with 100 μM DCF for different time points (0–72 h) in NKE cells. (E) The scratch assay was performed using different concentrations of DCF in HeLa cells for 24 and 48 h timepoints. (F) The relative fold change of the cell-free scratch area was estimated. (G) ROS levels were measured by H2DCFDA staining followed by FACS analysis. HeLa cells were treated with 175 μM of DCF at different timepoints. The bar diagram indicates the relative fold change in DCFDA fluorescence population compared with the control (no DCF treatment). (H) HeLa cells were treated with different concentrations of DCF and E7070 (100 μM) for 24 h. Cell cycle analysis was performed using PI staining followed by FACS analysis. The bar diagram depicts the relative fold change in the population of sub-G1 phase cells with different dosages of DCF. (I) Lipid peroxidation was assessed by measuring MDA levels by the TBARS method with different doses of DCF. MDA levels are represented as a fold change over control. ∗P<0.05, ∗∗P<0.01, ∗∗∗P<0.001, ns: non-significant. All data represent the mean ±SD of atleast three independent experiments.

To determine whether the cytotoxic effect rendered by DCF was cancer cell-specific, similar experiments were performed on two non-cancer epithelial cells (since the cancer cells used above were of epithelial origin): NKE cells (a normal kidney epithelial cell line) and HaCaT cells (skin keratinocytes). DCF did not significantly affect the viability of NKE cells at 175 μM (IC_50_ dose in HeLa cells), where the IC_50_ value was found to be greater than 1 mM concentration of DCF (Fig. 1C). Moreover, 100 μM DCF did not affect the viability of NKE cells at higher timepoints (48 and 72 h) (Fig. 1D) in comparison to HeLa cells (Fig. 1B). Similarly, in HaCaT cells, 175 μM DCF treatment did not induce a significant effect on the percentage cell viability, where the IC_50_ value of DCF was found to be greater than 1 mM concentration, suggesting the cytotoxic effects of DCF to be cancer-cell specific (Fig. S3).

To evaluate whether DCF inhibits directional cell motility, a scratch assay was performed with different concentrations of DCF over two time points (Fig. 1E). It was observed that DCF treatment inhibited directional cell motility in a dose- and time-dependent manner. The bar diagram (Fig. 1F) represents the relative fold change in the area of the scratch for different DCF concentrations and treatment time points that remained unoccupied by the cells.

To check whether DCF induces cell death in HeLa cells by elevating intracellular reactive oxygen species (ROS), a H2DCFDA staining assay followed by FACS analysis was performed at different time points after DCF treatment. The percentage distribution of DCFDA-positive cells was estimated when the DCF concentration was kept at 175 μM (Fig. S4). Results showed that 175 μM DCF significantly elevated intracellular ROS levels in a time-dependent manner, which confirmed that DCF induces oxidative stress in HeLa cells (Fig. 1G).

To check the effect of DCF on cell cycle progression, a PI staining assay was performed, followed by FACS analysis. HeLa cells were treated with different concentrations of DCF (87.5 and 175 μM) for 24 h, and E7070 was used as a positive control [26]. The percentage distribution of cells in the different phases of the cell cycle using different concentrations of DCF and E7070 was estimated from FACS analysis (Fig. S5A, S5B). 87.5 and 175 μM DCF treatment significantly increased the sub-G1 population in cells over the control cells, which is an indication of DNA damage (cell death) (Fig. 1H).

Excessive ROS levels lead to the lipid peroxidation of polyunsaturated fatty acids (PUFA), which is indicative of cellular oxidative stress [27]. Lipid peroxidation was estimated by measuring the levels of malondialdehyde (MDA) in HeLa cells. The result showed that DCF treatment in HeLa cells led to a significant increase in MDA levels, suggesting an increased level of lipid peroxidation (Fig. 1I).

### DCF induces apoptosis-mediated cell death in HeLa cells

To confirm whether DCF induces apoptosis in HeLa cells, Annexin-V/PI dual staining was performed with different concentrations of DCF for 24 h [27]. The percentage distribution of DCF-treated HeLa cells in different phases of apoptosis and necrosis was estimated by FACS analysis (Fig. S6). The data revealed that DCF treatment significantly elevated early and late apoptotic cell populations over control. However, there was no significant alteration of the necrotic population, which primarily indicates that DCF treatment induces apoptosis-mediated cell death (Fig. 2A, S6). Since apoptosis induces changes in the mitochondrial membrane potential (ΔΨm), we assessed whether DCF induced any changes in the ΔΨm by the JC-1 staining method. The percentage distribution of monomeric JC-1-positive (green fluorescence) HeLa cells with different doses of DCF was estimated by FACS analysis (Fig. 2B). DCF treatment increased the JC-1 (green) fluorescence dose-dependently after 24 h of incubation, which indicates the lowering of the mitochondrial membrane potential (Fig. 2B), another signature of apoptosis-mediated cell death [28].

**Fig. 2.**
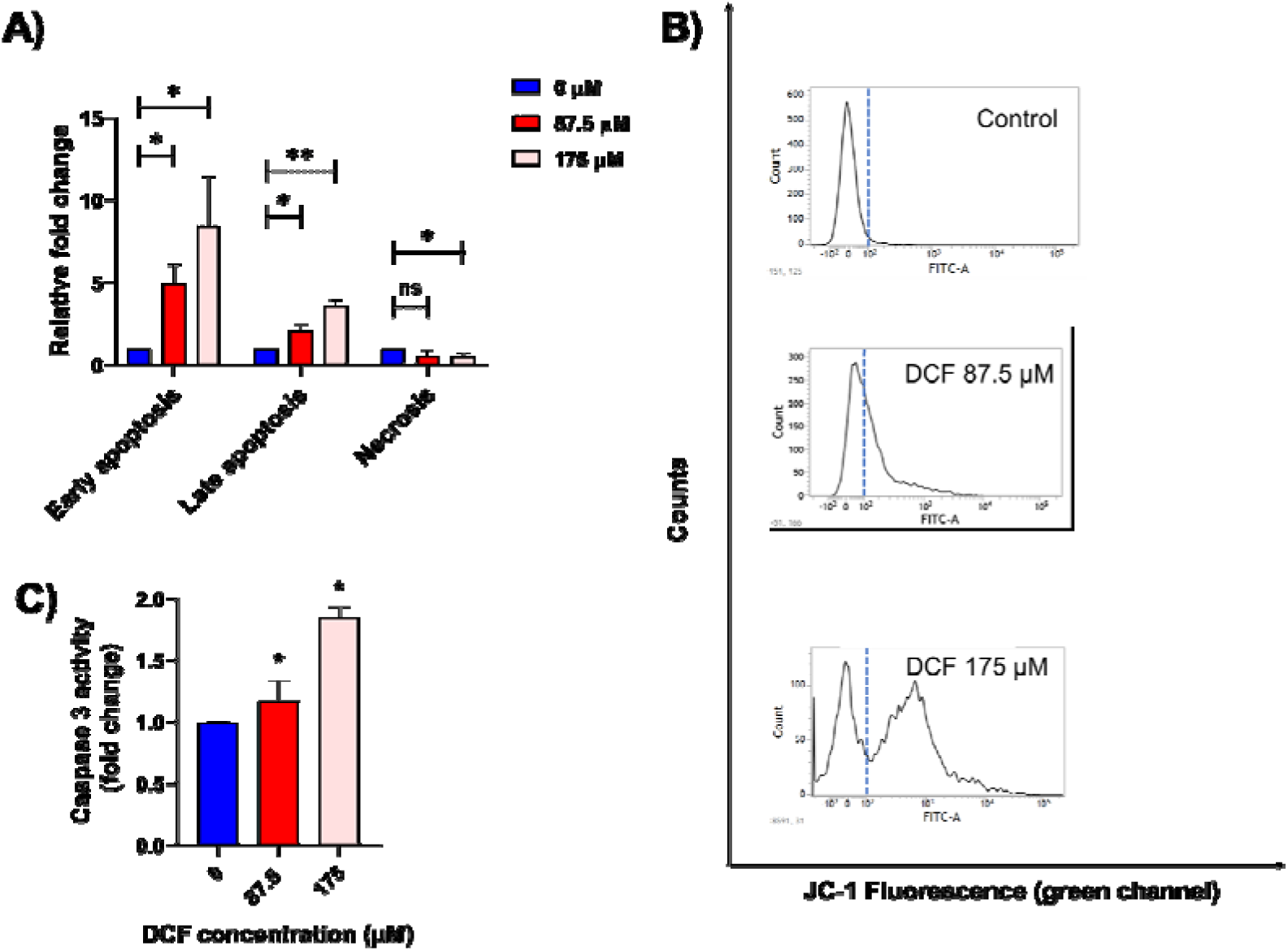
DCF induces apoptosis-mediated cell death in HeLa cells. (A) Annexin V/PI dual staining was performed, followed by FACS analysis using untreated and treated HeLa cells with different concentrations of DCF for 24 h. The bar diagram indicates the relative fold changes in the different phases of apoptotic and necrotic populations with DCF treatment. (B) Mitochondrial membrane potential (ΔΨm) was estimated in HeLa cells with different concentrations of DCF (24 h) by JC-1 staining followed by FACS analysis. (C) Caspase-3 activity in HeLa cells was estimated after treatment with different doses of DCF (24 h) and compared to the control. The bar graph represents the relative fold change of intracellular caspase-3 activity over control. All data represent the mean ± SD of at least three independent experiments. ∗P<0.05, ∗∗P<0.01, ns: non-significant compared to the control group.

Next, we assessed the effect of DCF on the activation of downstream caspase-3, one of the major effector caspases, whose activity is significantly increased during apoptosis [29]. Results showed that 24 h of DCF treatment significantly increased caspase-3 activity in Hela cells in a dose-dependent manner (Fig. 2C). All of these findings collectively imply that DCF causes apoptosis-mediated cell death in HeLa cells.

### DCF inhibits intracellular LDH-A in HeLa cells

Previous work shows that DCF demonstrates its antimitotic activity by interfering with the glucose metabolism route, where glycolysis is the primary glucose metabolism pathway in cancer cells [23]. Based on these findings, we hypothesized that some glycolytic enzymes may be potential DCF-binding partners at the cellular level. To determine this, molecular docking was performed using DCF with individual glycolytic enzymes. A higher binding score represents a more favorable binding. The result showed that the docking score for the LDH-A/DCF complex was the highest among other glycolytic enzyme-DCF complexes, which suggests that LDH-A could be a binding partner of DCF among the other glycolytic enzymes (Fig. 3A).

**Fig. 3.**
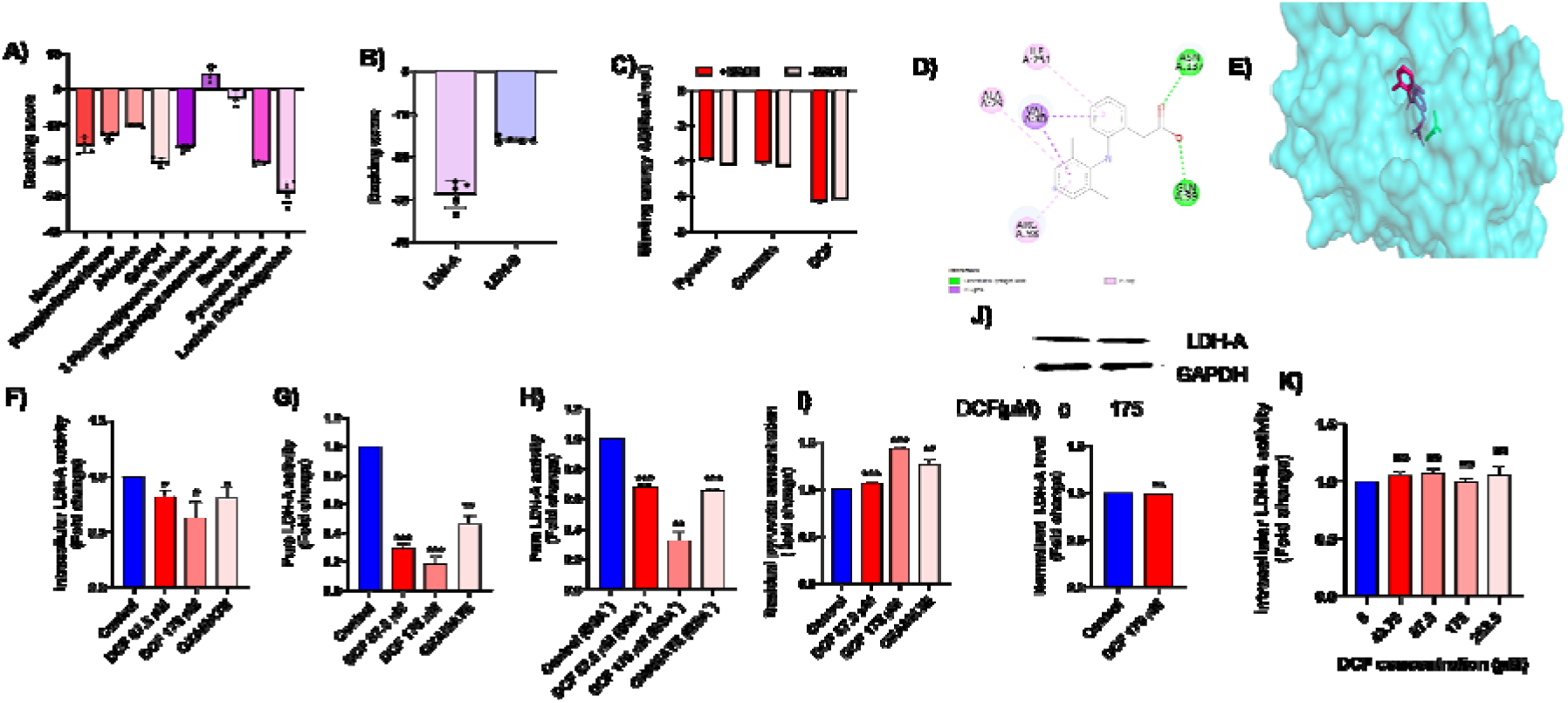
DCF inhibits intracellular LDH-A in HeLa cells. (A) Determination of the binding score of DCF with glycolytic enzymes. The crystal structures of the glycolytic enzymes were obtained from the PDB database. (B) Docking score of DCF with LDH-A and LDH-B. (C) Estimation of the binding energy (kcal/mol) of LDH-A with pyruvate (a substrate of LDH-A), oxamate (a known inhibitor of LDH-A), and DCF in the presence and absence of NADH. (D) 2D residue-level interaction of the LDH-A/DCF complex. Each dotted line indicates specific types of interaction mentioned in the figure. Interacting amino acids in LDH-A and their residue numbers have been specified. (E) 3D interaction of DCF and pyruvate with LDH-A. Red and blue-colored ligands indicate DCF, while green-colored ligands indicate pyruvate. (F) Intracellular LDH-A activity was measured with different concentrations of DCF in HeLa cells after 24 h of incubation with oxamate as a positive control. The bar diagram indicates the fold change in the enzymatic activity with respect to the control. (G) The effect of different concentrations of DCF on pure LDH-A activity. Oxamate is the positive control. The bar diagram indicates the relative fold change of average enzymatic activity over control. (H) Effect of DCF on LDH-A activity in the presence of 100 μM BSA (a crowding agent) was measured using DCF and oxamate. The bar diagram indicates the relative fold change of the average enzymatic activity over control. (I) The effect of DCF on pyruvate accumulation was estimated using DCF and oxamate. The bar diagram represents the relative fold change in the leftover pyruvate level after 3 minutes of the reaction. (J) Western blot analysis of LDH-A expression in HeLa cells after 175 μM DCF treatment for 24 h. Expression of GAPDH was used as a loading control. The bar diagram indicates the relative fold change in the normalized expression of LDH-A/ GAPDH levels due to DCF treatment. (K) *in-vitro* LDH-B activitity was estimated with the indicated concentrations of DCF at 24 h in HeLa cells. All the bar graphs represent the mean ± SD of three independent experiments where ∗P<0.05, ∗∗P<0.01, ∗∗∗P<0.001 and ns: non-significant compared to the control group.

To check the binding specificity of DCF towards LDH-A and LDH-B, *in-silico* binding scores were determined using molecular docking analysis. It was found that DCF has a higher binding score with LDH-A than with LDH-B, implying that LDH-A has a greater binding affinity with DCF than LDH-B (Fig. 3B).

An *in-silico* docking using AutoDock was performed to check the binding energy of DCF, oxamate (a known inhibitor of LDH-A), and pyruvate (a substrate of LDH-A) with LDH-A in the presence or absence of NADH. At first, AutoDock was validated by redocking oxamate with LDH-A and comparing the interacting residues with those of the crystal structure of LDH-A bound with oxamate. Results indicated that redocking of oxamate with LDH-A interacted with similar amino acids when compared to the database (Fig. S8) [30]. We found that DCF had a higher binding free energy than oxamate or pyruvate (Fig. 3C), and the presence of NADH did not cause any significant effect on DCF binding with LDH-A (Fig. 3C), as both binding energies in the presence and absence of NADH remained similar [12]. Docking studies also revealed that DCF interacted with LDH-A in the vicinity of the pyruvate binding site, predicting that DCF could be an allosteric inhibitor of LDH-A (Fig. 3D, 3E, S9).

To validate our *in-silico* predictions and to check whether DCF had any effect on LDH-A activity, intracellular LDH-A activity was examined using different concentrations of DCF in HeLa cells. The data demonstrated that DCF inhibited intracellular LDH-A activity dose-dependently (Fig. 3F). To determine whether DCF inhibited intracellular LDH-A activity in other cancer cell lines, intracellular LDH-A activity in both HCT-116 and MCF-7 cell lines was estimated using different doses of DCF, using oxamate as a positive control. It was observed that DCF dose-dependently inhibited intracellular LDH-A activity in both HCT-116 and MCF-7 cells, which suggested that the inhibitory action of DCF towards LDH-A is not cervical cancer cell-specific (Fig. S10, S11). To further establish whether LDH-A is a direct target of DCF, the same enzyme activity assay was performed with pure muscle LDH-A using different concentrations of DCF; oxamate was used as a positive control, and the absence of NADH was used as a negative control (Fig. S12). Results showed that DCF inhibited pure LDH-A activity in a dose-dependent manner (Fig. 3G). Moreover, DCF dose-dependently reduced pure LDH-A activity in the presence of 100 μM BSA (which served as a crowding agent) (Fig. 3H), revealing that DCF still found its binding specificity with LDH-A even in a molecularly crowded background [31]. We also measured the substrate (pyruvate) utilization by pure LDH-A with different concentrations of DCF. It was observed that pyruvate levels accumulated with increasing concentrations of DCF, suggesting that the accumulation of the substrate was due to the inhibition of the forward reaction (Fig. 3I).

To check whether DCF inhibits LDH-A at the translational or post-transcriptional level, western blot analysis was performed, and LDH-A expression was evaluated in the absence and presence of DCF (Fig. 3J). The data clearly reveals that DCF did not significantly alter LDH-A expression at the protein level compared to the control, which suggests that DCF inhibits LDH-A at the post-translational level.

To evaluate whether DCF has any effect on the other isoform of LDH, intracellular LDH-B activity was evaluated in HeLa cells with varied doses of DCF. The result demonstrates that DCF did not significantly affect intracellular LDH-B activity, indicating that the cellular effects brought about by DCF are specifically caused by the inhibition of the LDH-A isoform (Fig. 3K). Similarly, DCF treatment did not significantly affect intracellular LDH-B activity in the MCF-7 cell line (Fig. S13). This result correlates well with our *in-silico* prediction that shows DCF has a greater binding affinity with LDH-A compared to LDH-B (Fig. 3B).

### DCF affects enzyme kinetics by interacting with LDH-A

To determine the mode of inhibition of LDH-A by DCF, the reaction kinetics were monitored with different concentrations of pyruvate, keeping the NADH concentration constant in the presence and absence of DCF. In the absence of the inhibitor DCF, normal Michaelis-Menten enzyme kinetics were observed. However, in the presence of DCF, the enzyme kinetics showed a typical ‘J-shaped’ curve, which is a signature of allosteric inhibition (Fig. 4A) [32].

**Fig. 4.**
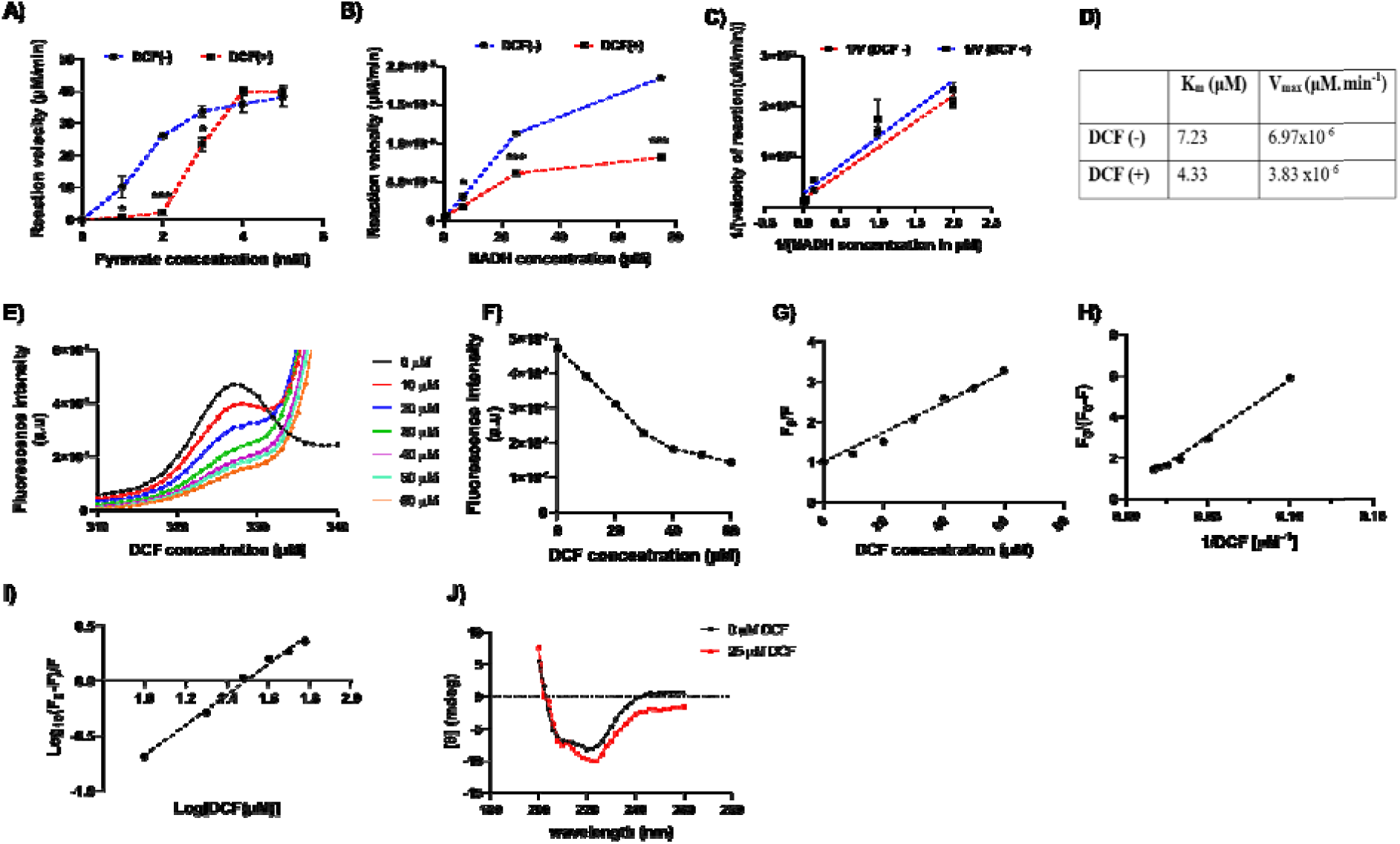
DCF alters enzyme kinetics by interacting with LDH-A. (A) Reaction kinetics of LDH-A with pyruvate (1–5 mM) were monitored in the presence and absence of DCF (175 μM) with a fixed NADH concentration (150 μM). (B) The reaction kinetics of LDH-A with NADH (0.5–75 μM) were monitored in the presence and absence of DCF, where the pyruvate concentration was kept constant (1 mM). (C) Line Weaver bark plot to determine the parameters of the reaction kinetics (K_m_ and V_max_) of LDH-A in the presence and absence of DCF treatment. (D) The values of K_m_ and V_max_ in the presence and absence of DCF treatment are indicated. (E) Effect of DCF on the intrinsic fluorescence spectra of LDH-A with different concentrations of DCF (0–60 μM). The excitation wavelength was fixed at 295 nm. (F) Effect of DCF on the maximum fluorescence intensities obtained from fluorescence quenching data. (G) SV plot of the LDH-A/DCF complex at different concentrations of DCF, the quenching constant K_q_ was determined from the fluorescence quenching data. (H) Determination of the binding constant (K_a_) of the DCF/LDH-A complex obtained from a modified SV plot. (I) The double-logarithmic plot used to determine the number of DCF binding sites (n) on LDH-A. (J) Effect of DCF on the secondary structure of LDH-A using far-UV CD spectroscopy (200–260 nm).

To determine the effect of the cofactor NADH on DCF-induced inhibition of LDH-A, reaction kinetics were estimated with different concentrations of NADH while keeping the pyruvate concentration constant (Fig. 4B). The Michaelis-Menten curve and Lineweaver Burk plot (Fig. 4B, 4C) indicated an uncompetitive mode of inhibition since both K_m_ and V_max_ values were altered (Fig. 4D) [33]. To determine the binding constant for the LDH-A/DCF complex, the effect of DCF on the intrinsic tryptophan fluorescence of LDH-A was determined [34, 35]. DCF inhibited the intrinsic tryptophan fluorescence of LDH-A in a dose-dependent manner, showing that DCF interacts with LDH-A and induces structural changes (Fig. 4E, 4F).

For determining the quenching constant (K_q_) of the DCF/LDH-A complex, the Stern-Volmer (SV) equation was used [36]

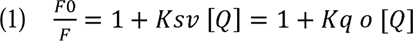

Where F and F_0_ are the fluorescence intensities in the presence and absence of DCF, [Q] is the quencher concentration (DCF) in μM, K_sv_ is the Stern-Volmer constant that estimates the efficiency of quenching, K_q_ is the quenching rate constant of the protein, a is the average lifetime of the fluorophore (tryptophan) in the absence of the quencher, which is roughly 5.7 ns [37]. From fluorescence quenching data (Fig. 4F,4G) the value of K_sv_ was calculated to be 3.71 x 10^4^ L. mole^-1^. This K_sv_ was used to calculate the quenching rate constant K_q_. Reports suggest that if the K_q_ value is greater than 2 x 10^10^ L.mol^-1^.sec^-1^ the quenching process is static, and if it is smaller, the quenching is dynamic [37]. The K_q_ value obtained from our study (equation 1) was 6.49 x 10^12^ L.mol^-1^.sec^-1^ which indicates that the quenching was mainly static in nature (Fig. 4G). So, DCF formed a ground-state complex with LDH-A. For the static quenching process, this fluorescence quenching data can be further analyzed by using the modified Stern-Volmer equation [38].

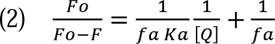

Where f_a_ is the mole fraction of the solvent-accessible fluorophore, and K_a_ is the binding constant. From fitting the fluorescence quenching data (Fig. 4H) into the equation (2) K_a_ obtained was 0.00593 μM^-1^

The binding constant (K_a_) and the binding site number (n) can also be analyzed from the double logarithmic equation [37]

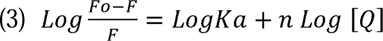

By fitting the fluorescence quenching data into the equation (3), the binding constant (K_a_) was obtained as 0.008703 μM^-1^ (Fig. 4I), which is equivalent to the binding constant value obtained from the previous equation (2) (Fig. 4H). The number of DCF binding sites on LDH-A was found to be approximately 1 (n=1.38).

The effect of DCF (25 μM) on the secondary structure of LDH-A was monitored by far-UV CD spectroscopy [36, 39]. The results revealed that the interaction of DCF causes a considerable alteration in the secondary structure of LDH-A (Fig. 4J, S14). On DCF binding, the percentage of alpha helix changed from 82% to 45.8%; beta sheets changed from 5.7% to 34.4%; and turn changed from 12.2% to 14.9% (Fig. S14).

### DCF alters the level of intracellular metabolites via the inhibition of LDH-A

The effect of LDH-A inhibition on intracellular pyruvate level, extracellular acidification, NAD^+^/NADH ratio, and glycolytic flux in HeLa cells was studied at two separate time periods (24 h and 48 h) with different concentrations of DCF. It was found that DCF treatment led to an increase in the intracellular pyruvate concentration in a dose- and time-dependent manner (Fig. 5A). Likewise, DCF increased intracellular pyruvate levels in MCF-7 cells in a dose-dependent manner. (Fig. S15).

**Fig. 5.**
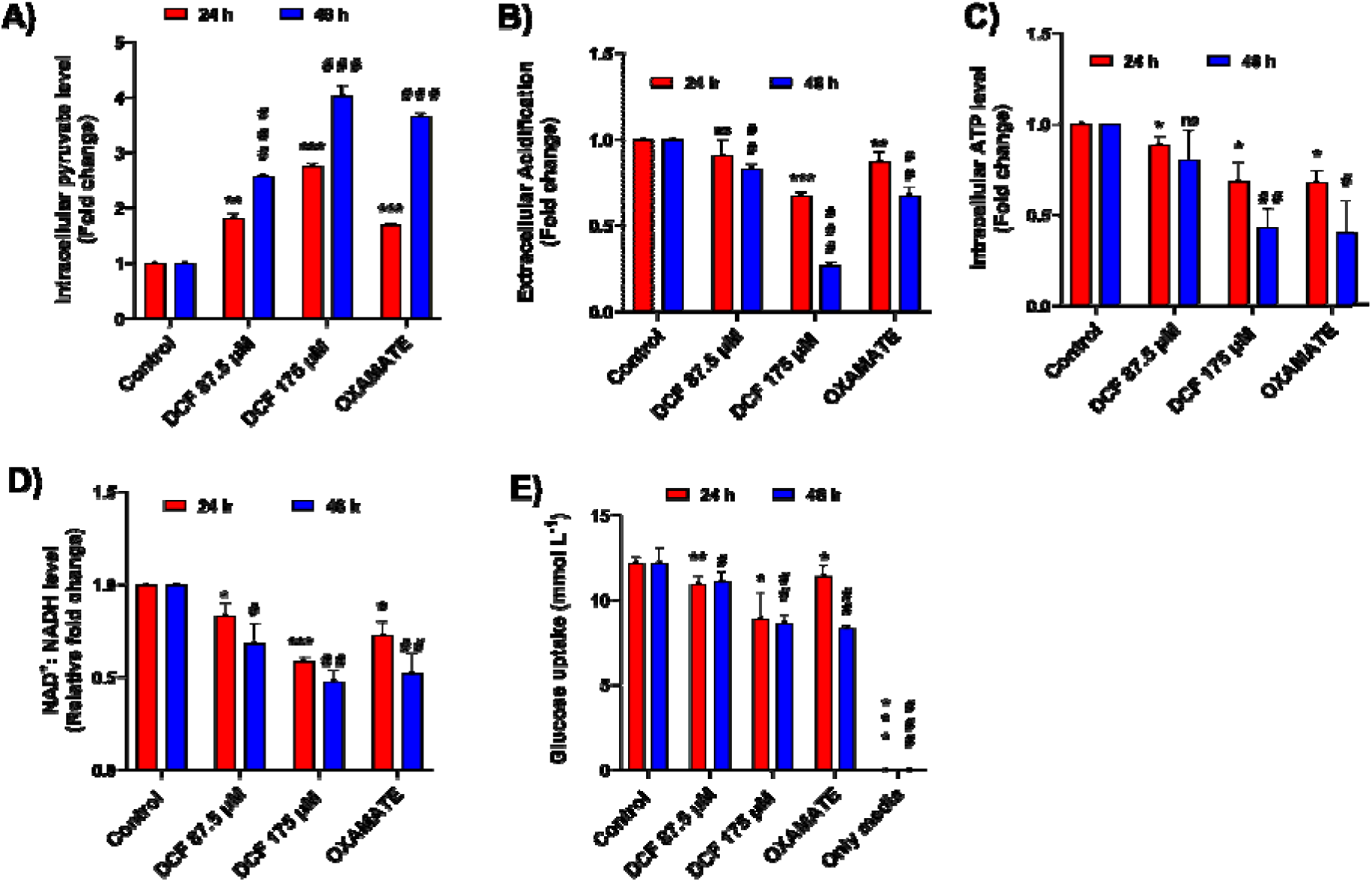
DCF alters intracellular metabolite levels via inhibition of LDH-A. Effect of DCF on (A) intracellular pyruvate level; (B) extracellular acidification; (C) intracellular ATP level; (D) intracellular NAD^+^/NADH ratio; and (E) glucose uptake at 24 h and 48 h time points. Oxamate was used as a positive control. The bar graph represents the mean ± SD of three independent experiments. ∗P<0.05, ∗∗P<0.01, and ∗∗∗P<0.001 compared to the control group for 24 h. #P<0.05, ##P<0.01, and ###P<0.001 compared to the control group at 48 h.

On the other hand, DCF dose- and time-dependently lowered the extracellular acidification rate in HeLa cells (Fig. 5B). This alteration in extracellular acidification by DCF was confirmed by the alteration of the extracellular pH level in HeLa cells (Fig. S16). Likewise, DCF also lowered extracellular acidification in MCF-7 and HCT-116 cells in a dose-dependent manner (Fig.S18 A, B). This lowering of extracellular acidification by DCF in turn reduced the glycolytic flux in HeLa cells (Fig. 5E) as well as in MCF-7 and HCT-116 cells (Fig. S20). For the extracellular acidification and the glucose uptake assays, oxamate was used as a positive control, whereas no cell, no cell +DCF, and no cell +oxamate served as negative controls (Fig. S17A, B).

Intracellular ATP levels were estimated using different doses of DCF, and it was observed that DCF reduced intracellular ATP levels dose- and time-dependently in HeLa cells (Fig. 5C). Similarly, DCF dose-dependently lowered intracellular ATP levels in MCF-7 cells (Fig. S19). The effect on intracellular NAD^+^/NADH ratio was also checked with different concentrations of DCF at 24 and 48 h in HeLa cells. It was observed that DCF lowered the NAD^+^/NADH ratio in a time- and dose-dependent manner (Fig. 5D). Oxamate was used as a positive control for all experiments. These results confirmed the inhibition of LDH-A activity by DCF at the cellular level.

### DCF causes anti-mitotic effects by specifically inhibiting LDH-A activity in HeLa cells

siRNA-mediated specific knockdown of LDH-A was carried out to assess the selectivity of DCF towards LDH-A in HeLa cells. si-LDH-A knocked down LDH-A successfully after 48 h (Fig. 6 A, B). This si-LDH-A-mediated knockdown of LDH-A in HeLa cells led to a significant decrease in percentage cell viability (Fig. 6C), intracellular LDH-A activity (Fig. 6D), extracellular acidification (Fig. 6E), glucose uptake (Fig. 6F), and intracellular ATP levels (Fig. 6G). These findings are consistent with those obtained from DCF-treated HeLa cells (Fig. 1A, 3F, 5B, 5C, 5E). Interestingly, 24 h of incubation with varied doses of DCF did not cause any significant differences in cell viability or extracellular acidification in si-LDH-A transfected cells when compared to control scrambled siRNA transfected cells or non-transfected cells, which showed a dose-dependent decrease in cell viability and extracellular acidification (Fig. 6H, I). The cells lost their sensitivity towards DCF in the absence of LDH-A, confirming that DCF renders its anti-mitotic effects via LDH-A inhibition.

**Fig. 6.**
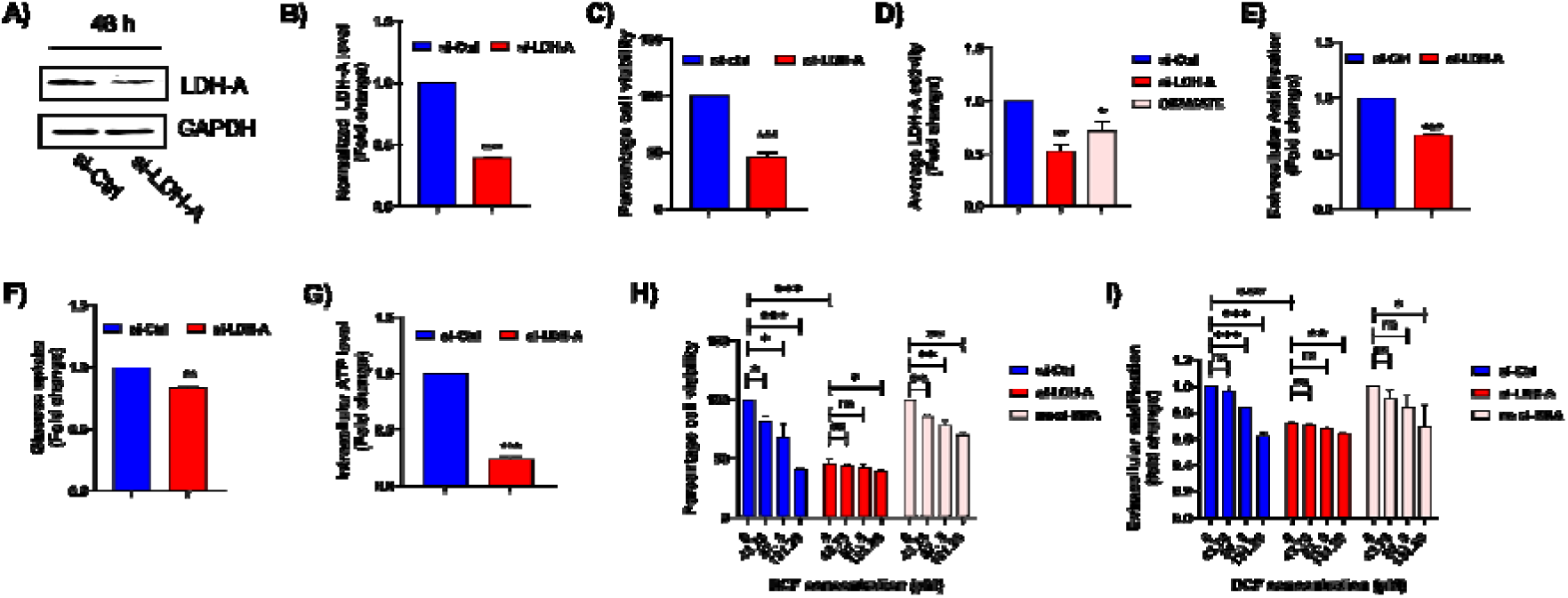
DCF specifically inhibits LDH-A activity in HeLa cells. (A) HeLa cells were transfected with control siRNA and si-LDH-A for 48 h, and LDH-A expression was determined by western blot analysis. (B) The bar diagram indicates the relative fold change in the normalized expression of LDH-A/GAPDH levels in si-LDH-A transfected HeLa cells compared to control scrambled siRNA transfected cells. Determination of the effect of si-LDH-A transfected HeLa cells on (C) percentage cell viability; (D) average fold change in intracellular LDH-A activity; (E) average fold change in extracellular acidification; (F) average fold change in glucose uptake; and (G) average fold change in intracellular ATP level in comparison with control siRNA transfected cells at 48 h. Effect of different concentrations of DCF on H, cell viability (by trypan blue exclusion assay), and I, extracellular acidification after 48 h of transfection of si-LDH-A, control siRNA, and non-transfected Hela cells. The bar graph represents the mean ± SD of three independent experiments. ∗P<0.05, ∗∗P<0.01, and ∗∗∗P<0.001, ns P> .05, ns: non-significant as compared to the control group.

### DCF inhibits hypoxia in HeLa cells

Cobalt chloride (CoCl_2_) has been widely used for establishing hypoxic microenvironments in a variety of cancer cell lines, even in the presence of sufficient oxygen conditions, and LDH-A is a key enzyme that is overexpressed in the hypoxic milieu [40, 41]. To induce LDH-A expression in HeLa cells, cells were treated with different concentrations of CoCl_2_ (0–100 μM) for 24 h. CoCl_2_ treatment caused an increase in cell viability in a dose-dependent manner, where 100 μM CoCl_2_ concentration was responsible for a significant increase in cell viability (Fig. 7A). Moreover, 100 μM CoCl_2_ resulted in a significant increase in extracellular acidification (Fig. 7B) as well as LDH-A expression (Fig. 7C, D). Since LDH-A is a marker of hypoxia and LDH-A expression is maximal at 100 μM CoCl_2_ concentration, we chose this CoCl_2_ concentration to induce LDH-A expression for our further experiments.

**Fig. 7.**
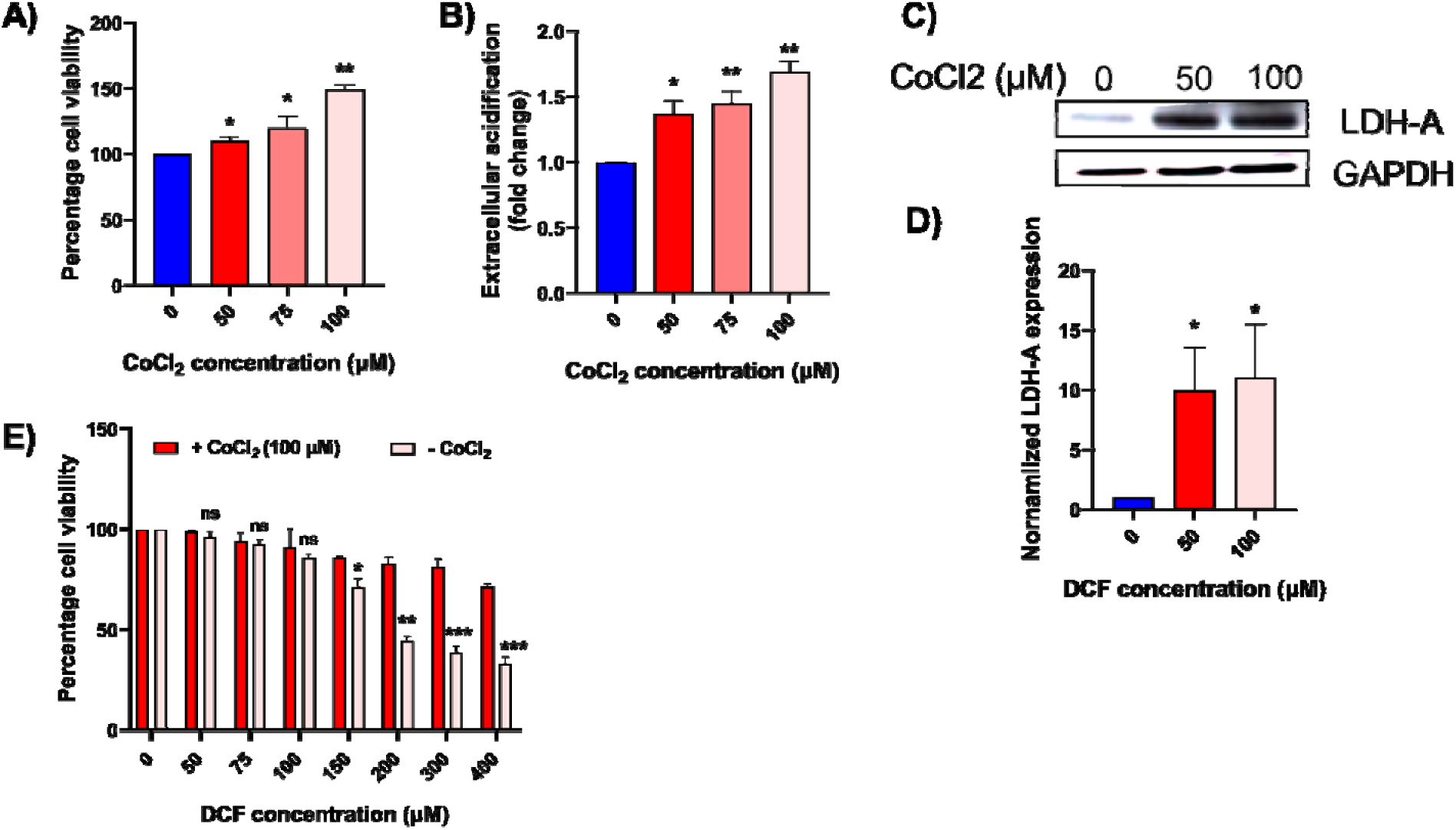
CoCl_2_-induced hypoxia decreases the sensitivity of HeLa cells to DCF. (A) HeLa cells were treated with the indicated concentration of CoCl_2_ (0-100μM) for 24 h, and cell viability was estimated by trypan blue exclusion assay. (B) Extracellular acidification was estimated in HeLa cells with different concentrations of CoCl_2_ (0–100 μM) at 24 h. (C) Western blot analysis of LDH-A expression in HeLa cells after CoCl_2_ (0,50,100 μM) treatment for 24 h. GAPDH was used as a loading control. (D) The bar diagram represents the relative fold change in the normalized expression of LDH-A/GAPDH levels with different concentrations of CoCl_2_. (E) Effect of varying DCF concentrations (24 h incubation) on Hela cell viability in the presence and absence of CoCl_2_ (100 μM)-induced hypoxia. The bar graph represents the mean ± SD of three independent experiments. ∗P<0.05, ∗∗P<0.01, and ∗∗∗P<0.001, ns P> .05, ns: non-significant as compared to the control group.

To evaluate whether HeLa cells are sensitive to DCF by overexpressing a hypoxic microenvironment, we estimated cell viability in the presence and absence of 100 μM CoCl_2_ at 24 h. The result showed that cells treated with 100 μM CoCl_2_ lost their sensitivity towards DCF in comparison with untreated cells since higher DCF concentrations were required for rendering cytotoxic effects. Moreover, the IC_50_ value of DCF was higher in CoCl_2_-induced hypoxic HeLa cells (more than 400 μM) compared to the control (no CoCl_2_ treatment) (175 μM) (Fig. 7E). This result proposes that DCF induces cell death by inhibiting the hypoxic microenvironment in HeLa cells via inhibition of LDH-A [42].

### DCF induces cell death via inhibition of LDH-A and activation of the AMPK signaling pathway

Our findings showed that DCF triggers apoptosis-mediated cell death in HeLa cells in a dose- and time-dependent manner via decreasing intracellular LDH-A activity (Fig. 2A–C, 3F, 3G, 6H). We also showed that DCF-induced suppression of LDH-A activity reduced intracellular ATP levels (Fig. 5C). Since AMPK is sensitive to changes in the intracellular AMP/ATP ratio, we investigated the phosphorylation of AMPK in the presence and absence of DCF. The results showed that DCF treatment increased AMPK phosphorylation compared to the control (Fig. 8A, B). As AMPK activation induces apoptosis by inhibiting its downstream target, p-70S6K [43], we assessed the phosphorylation of p-70S6K by DCF treatment and found that DCF treatment suppressed the phosphorylation of S6K (Fig. 8A, C).

**Fig. 8.**
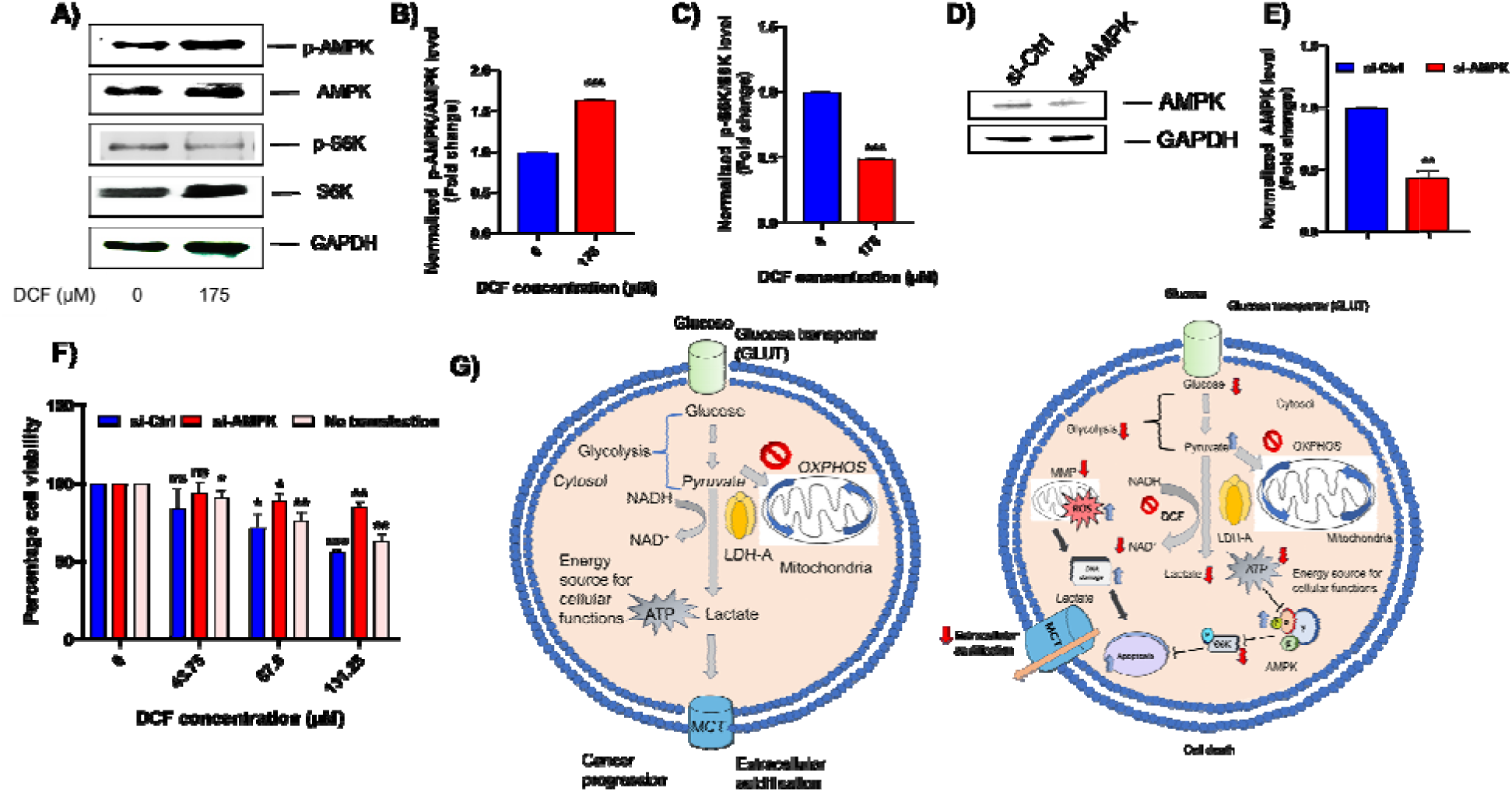
DCF induces cell death via activation of the AMPK signaling pathway. (A) Western blot analysis of p-AMPK (Thr-172), AMPK, p-S6K, and S6K expression in HeLa cells after 175 μM DCF treatment. (B) Representative bar graph of the relative fold change of the normalized expression of p-AMPK/AMPK and (C) p-S6K/S6K in DCF-treated HeLa cells compared to the control (no DCF). (D) AMPK, and GAPDH expression in HeLa cells transfected with si-Ctrl and si-AMPK transfection for 48 h. (E) Relative fold change of the normalized expression of p-AMPK/GAPDH in si-Ctrl and si-AMPK transfected cells at 48 h. (F) Effect of different concentrations of DCF on the percentage cell viability in si-Ctrl, si-AMPK, and non-transfected HeLa cells using trypan blue exclusion assay. siRNA transfection was carried out at 48 h, followed by DCF treatment for 24 h. The bar graph represents the mean ± SD of three independent experiments. ∗P<0.05, ∗∗P<0.01, and ∗∗∗P<0.001, ns P> .05, ns: non-significant as compared to the control group. (G) Model for the mode of action of DCF that leads to apoptosis in HeLa cells.

To evaluate whether DCF induces cell death in HeLa cells by activation of the AMPK pathway, knockdown of AMPK by siRNA transfection was performed in HeLa cells. AMPK-specific siRNA significantly reduced the expression of AMPK in HeLa cells at 48 h. GAPDH expression was used as a loading control (Fig. 8 D, E). When percentage cell viability was estimated with different concentrations of DCF in si-AMPK, control siRNA (si-Ctrl), and non-transfected HeLa cells at 24 h, DCF did not significantly affect cell viability in si-AMPK-transfected HeLa cells in comparison with control siRNA and non-transfected HeLa cells, confirming the involvement of AMPK in DCF-induced cytotoxicity (Fig. 8 F). Various supporting data showed that LDH-A inhibition activates AMPK in cancer cells [44, 45]. In our study, to correlate the involvement of LDH-A inhibition with AMPK activation (sensitive to intracellular ATP fluctuations), we found that intracellular ATP levels were depleted via LDH-A inhibition (by si-LDH-A), suggesting the involvement of LDH-A in AMPK activation (Fig. 6G). This was checked by assessing p-S6K levels (the downstream target of activated AMPK) when intracellular LDH-A was knocked down by si-LDH-A transfection in HeLa cells. The results showed that knockdown of LDH-A inhibited p-S6K (Fig. S21). Since both siRNA-mediated knockdown of LDH-A and DCF-mediated inhibition of LDH-A depleted intracellular ATP levels and lowered S6K activation, we could ascertain the involvement of LDH-A inhibition by DCF in the AMPK signaling pathway. As a result, we infer that DCF triggers cell death in HeLa cells by targeting LDH-A, which decreases intracellular ATP levels, activates AMPK, and induces apoptosis via inhibiting S6K activation (Fig. 8G).

## Discussion

Cancer cells exhibit varying degrees of enhanced glycolytic rate depending on their cell types and growth preferences. Thus, ‘addiction to glycolysis’ and cancer progression share a very strong correlation [1, 2, 3]. Even when sufficient oxygen is available, cancer cells prefer glycolytic fermentation over mitochondrial respiration due to a hypoxic microenvironment and immediate energy demand [3, 43]. A cancer cell benefits from metabolic reprogramming not just in terms of rapid energy generation but also in terms of extracellular acidification of the tumor microenvironment, which has been identified as one of the key hallmarks of cancer [4]. Targeting cancer metabolism to restrict tumor progression is poorly understood and still remains a black box because of the altered metabolic preferences of different types of cancer cells. Thus, the search for novel drugs to arrest these metabolic switches in cancer cells is a global challenge for scientists. LDH-A serves as a key enzyme for metabolic reprogramming, instant energy production, and maintaining tumor homeostasis in cancer cells [46]. As LDH-A is overexpressed and hyperactive in many cancers, a variety of reports suggest that inhibiting LDH-A leads to cell death, thus conferring the dependency of cancer cells on LDH-A [11, 12, 13].

In this study, we show that DCF, an NSAID, causes apoptosis-mediated cell death in HeLa cells (IC_50_ dose 175 µM) and other cancer cells (Fig.1A-B, S1, S2, 2A-C). Since the profiles for glucose metabolism between normal cells and cancer cells are very different [47, 48], choosing control cervical cell lines or other control cell lines would not serve as an appropriate control for our study. In the two control normal cells, NKE and HaCaT, LDH-A activity was significantly lower compared to the HeLa cells (Fig. S7). Also, previous work showed that in normal cervical cells, LDH-A expression levels were significantly lower than in cervical cancer cells [13, 49]. However, in normal cells (NKE and HaCaT), we find that the dose of DCF chosen for this study (175 µM, IC_50_ in HeLa cells) causes only 10% cell death compared to cancer cells (Fig. 1C, S3). This suggests that the cytotoxic effect of DCF is cancer cell-specific. Additionally, we show that LDH-A is a target of DCF. DCF binds to LDH-A, as confirmed from *in-silico* and *in-vitro* studies (Fig. 3A-I, 4A, 4B) and biophysical studies that revealed DCF-induced structural changes in LDH-A (Fig. 4E, 4J, S14). DCF binding did not affect LDH-A expression at the protein level (Fig. 3J) [50], but post-translationally inhibited LDH-A activity both in a cell-free system (Fig. 3G, 3H, 3I) and in cells (Fig. 3F, S10, S11), which establishes LDH-A as a probable target of DCF. DCF was found to allosterically inhibit pyruvate binding to LDH-A (Fig. 4A), which validates our *in-silico* prediction where we have shown that DCF binds with LDH-A in the vicinity of its substrate binding site by inducing a conformational change in the enzyme (Fig. 3D, 3E, S9).

Our *in-silico* prediction showed that the presence of the co-factor NADH did not significantly affect DCF binding with LDH-A, probably due to their different binding sites and comparable binding energies (Fig. 3C) [12]. Validating our *in-silico* results in a cell-free system, we found that DCF binding did not have any inhibitory effect on NADH binding to the enzyme, and the signature graph for the change in K_m_ and V_max_ values in the presence and absence of DCF suggested an uncompetitive mode of inhibition by DCF (Fig. 4B-D) [51].

LDH-A is a key enzyme that is responsible for the conversion of pyruvate to lactate in the glycolytic pathway [6, 7]. This forward reaction facilitates cancer cells by increasing glucose uptake, extracellular acidification, and intracellular ATP levels [52]. Many reports have suggested that inhibition of LDH-A is responsible for lowering the glycolytic flux [52, 53]. Similarly, DCF-induced inhibition of cellular LDH-A was found to reduce the glycolytic flux by lowering glucose uptake (Fig. 5E, S20), extracellular acidification (Fig. 5B, S18), and intracellular ATP levels (energy deprivation) (Fig. 5C, S19). This DCF-induced reduction in glycolytic flux caused cell death in cancer cells. The involvement of LDH-A in DCF-induced HeLa cell cytotoxicity was further confirmed by assessing cell viability in LDH-A knocked-down cells using LDH-A-specific siRNA. DCF treatment did not cause any significant differences in cell viability or extracellular acidification in si-LDH-A transfected cells when compared to control siRNA transfected cells or non-transfected cells, which showed a dose-dependent decrease in cell viability and extracellular acidification (Fig. 6 H, I). The LDH-A knocked-down cells lost their sensitivity to DCF. These results suggest that LDH-A is a specific target of DCF, and in its absence, DCF fails to induce dose-dependent cytotoxicity in HeLa cells, confirming that DCF renders its anti-mitotic effects via LDH-A inhibition rather than other glycolytic enzymes [54]. To check the effect of DCF on the other isozyme of LDH, an intracellular LDH-B activity assay was performed in both HeLa and MCF-7 cells with different concentrations of DCF (Fig. 3K, S13). Results suggest that DCF did not have any significant inhibitory effect on intracellular LDH-B activity, which also validates our *in-silico* finding that DCF showed lower binding affinity towards LDH-B than LDH-A (Fig. 3B).

Many reports suggest that there is a strong correlation between the inhibition of LDH-A and the induction of oxidative stress (ROS production) via mitochondrial damage [55, 56, 12]. According to Le et al., si-LDH-A transfection-mediated suppression of LDH-A increases ROS levels (DCFDA staining), which is responsible for oxidative stress in cancer cells [55]. Kim et al. report that PSTMB (an LDH-A inhibitor) induces apoptosis-mediated cell death by elevating mitochondrial ROS production in the HT29 cell line (12). This increase in ROS level is responsible for alteration of the cell cycle phases, lipid peroxidation, and lowering of mitochondrial membrane potential, which further initiates apoptosis-mediated cell death [27, 57, 58]. Assessing the effect of DCF on cellular ROS levels, we likewise found that inhibition of intracellular LDH-A by DCF (Fig. 3F) led to an increase in cellular ROS levels (Fig. 1G, S4), lipid peroxidation (Fig. 1I), cell cycle alteration (Fig. 1H, S5), change in mitochondrial membrane potential (Fig. 2B), and other apoptotic events in HeLa cells, including downstream caspase 3 activation (Fig. 2A, S6, 2C).

AMPK is a heterotrimeric sensor kinase that gets activated by the increasing concentration of AMP and ADP and the decreasing concentration of ATP [14, 15]. Activation of AMPK results in the phosphorylation of Thr-172 on its catalytic α subunit. This phosphorylation of AMPK occurs either by binding ADP or AMP to it [59]. AMP helps in maintaining the sustained activation of AMPK by inhibiting dephosphorylation, whereas ATP inhibits active AMPK by initiating dephosphorylation. Depletion of intracellular ATP, which activates AMPK activity, has been shown to inhibit cancer cell growth. Thus, AMPK plays a pivotal role in maintaining cell survival and apoptosis and serves as a potent target to treat a wide range of cancers [17, 18, 19]. Among them, an AMPK activator, 5-amino-1-β-D-ribofuranosyl-imidazole-4-carboxamide (AICAR), has been used to inhibit cancer growth [60]. Various natural compounds like resveratrol, quercetin, and 24-hydroxylursolic acid also induce apoptosis by activating AMPK [61, 62, 63]. Moreover, inhibition of LDH-A activity leads to glucose deprivation and energy starvation, which trigger apoptosis. LDH-A inhibition is strongly associated with AMPK activation. Several studies have demonstrated that the knockdown of LDH-A stimulates AMPK signaling in several cancer cell lines and induces apoptosis-mediated cell death [19, 44]. From our results, we propose that DCF (an LDH-A inhibitor) activates AMPK phosphorylation via depletion of intracellular ATP levels in HeLa cells (Fig 3F, 5C, 8A, 8B). This DCF-mediated AMPK activation further downregulates the p-S6K level, which in turn is responsible for DCF-mediated cell death (Fig. 8A, 8C). Furthermore, si-AMPK-mediated AMPK knockdown inhibited DCF-mediated cell death (Fig. 8F), confirming the activation and participation of the AMPK signaling pathway in DCF-induced apoptosis in HeLa cells [64]. To correlate the involvement of LDH-A inhibition with AMPK activation (sensitive to intracellular ATP fluctuations), we found that intracellular ATP levels were depleted in cells transfected with si-LDH-A, which knocked down LDH-A, justifying the involvement of LDH-A in AMPK activation (Fig. 6G). This was confirmed by inhibition of p-S6K levels (a downstream target of activated AMPK) in LDH-A knocked-down HeLa cells (Fig. S21).

Since both siRNA-mediated knockdown of LDH-A and DCF-mediated inhibition of LDH-A depleted intracellular ATP levels and lowered S6K activation, we could justify the involvement of LDH-A inhibition by DCF in the AMPK signaling pathway. However, the direct activation of the AMPK pathway by DCF (independent of LDH-A inhibition) cannot be ruled out based on our experiments.

As previously noted, hypoxia induction is a significant characteristic of cancer cells. Many investigations have demonstrated that hypoxia is strongly associated with the upregulation of intracellular LDH-A [65]. According to many studies, inhibiting LDH-A attenuates hypoxia in various types of cancer cell lines, thus decreasing cell proliferation and homeostasis due to the prevention of metabolic switching [42, 66]. In this study, inducing hypoxia in HeLa cells by CoCl_2_ treatment increased LDH-A expression and made the cells less sensitive to DCF-induced cytotoxicity compared to control cells (Fig. 7A-E). This implies that DCF-induced LDH-A inhibition prevents HeLa cells from exhibiting metabolic switching, which is responsible for cell death.

In conclusion, our work for the first time reports DCF, an NSAID group of drugs, as a specific LDH-A inhibitor that induces apoptosis in HeLa cells by activation of the AMPK pathway, mainly due to ATP deprivation in cells. This activated AMPK further reduces p-S6K expression, inhibiting cancer cell proliferation and progression. Thus, we have identified DCF as a lead anti-cancer molecule. Synthesizing more potent DCF derivatives for inhibiting LDH-A would enable optimal drug development to treat a wide range of "glycolytic-addicted" cancers.

## Experimental procedures

### Materials

Diclofenac (sodium salt), E7070, sodium oxamate, sodium pyruvate, 2’-7’-Dichlorodihydrofluorescein diacetate (H2DCFDA), 5,5,6,6’-tetrachloro-1,1’,3,3’ tetraethylbenzimi-dazoylcarbocyanine iodide (JC-1) was purchased from Sigma Aldrich; NADH, cobalt chloride (CoCl_2_) was purchased from SRL chemicals, India. Trypan Blue, Annexin-V binding buffer, Annexin V-FITC, and RNAiMAX were purchased from Thermo Fisher Scientific USA; Dulbecco’s modified eagle medium (DMEM), fetal bovine serum (FBS), and Opti-MEM media from Gibco; penicillin-streptomycin, trypsin-EDTA solutions, and propidium iodide were purchased from HiMedia, India. All the antibodies used in the experiments were purchased from Elabscience and Bio Bharati Life Science, India. Scrambled siRNA was purchased from Santa Cruz Biotechnology, USA, and specific siRNA from GenScript, USA.

### Cell culture

HeLa, HCT-116, MCF-7, and HaCaT cell lines were obtained from the National Centre for Cell Sciences Pune (NCCS). We obtained the NKE cell line as a gift from Prof. Kaushik Biswas’s laboratory, Bose Institute, India [67]. Cell lines were cultured in DMEM supplemented with 10% FBS, 1% antibiotics, 1% amphotericin, and 1% L-glutamine. Incubation was carried out at 37 °C in a 5% CO_2_ atmosphere.

### Cell viability assay

A trypan blue exclusion assay was carried out to determine the anti-mitotic effect of DCF. Cells were seeded at a density of 0.05 × 10^6^ (for HeLa, MCF-7, and HCT-116) and 1×10^5^ (for NKE and HaCaT) per well in a 24-well plate and incubated at 37°C. After 24 h, cells were treated with different concentrations of DCF. Then cells were collected, and trypan blue solution (0.4%) was added to the cell suspensions in a 1:1 ratio. Cells were counted using a hemocytometer. The percentage of cell viability was calculated and plotted.

### Scratch assay

HeLa cells were seeded at a density of 0.05 × 10^6^ cells per well in a 24-well plate to grow in a monolayer. After 24 h, a scratch was introduced in the confluent cell monolayer by a sterile pipette tip (20–200 μl). The detached cells were washed with PBS. After washing, 500 μL of fresh media was added, cells were treated with DCF to a final concentration of 87.5 and 175 μM, and the cell-free scratch area was quantified. Scratch zones were visualized under a microscope and photographed at 24 and 48 h.

### Measurement of ROS levels

ROS levels were evaluated by H2DCFHDA staining followed by FACS analysis. HeLa cells were seeded at a density of 4 ×10^5^ cells per well in a 6-well plate and incubated for 24 h. Then cells were treated with 175 μM of DCF for different timepoints and stained with 20 μM H2DCFDA in a serum-free medium for 30 minutes at 37°C. FACS Verse (BD FACS Verse, Bose Institute) was used to quantify DCFDA fluorescence, which is an indicator of intracellular ROS level [68].

### Cell cycle analysis using PI staining

Hela cells were plated in 6-well plates at a density of 0.3 x 10^6^ in each well. After 24 h, treatments were carried out with different concentrations of DCF and 100 μM of E7070 and incubated for 24 h. Cell pellets were resuspended in PBS. 40 μl ice-cold methanol was added for fixation. RNase A was added (final concentration 100 μg/μl) and the samples were kept in a 37°C water bath for 1.5 h. Propidium iodide (PI) was added (final concentration 120 μg/μl) for staining cellular DNA. FACS Verse was used to analyze the percentages of cells in different stages of the cell cycle by collecting at least 10,000 cells per sample [68].

### Measurement of lipid peroxidation

HeLa cells were seeded at a density of 4 x10^5^ per well in 6-well plates and incubated for 24 h. Then cells were treated with different concentrations of DCF for 24 h. Cells were collected and lysed after treatment by sonicating for 10 seconds. The thiobarbituric acid-reactive substances (TBARS) technique was used to analyze lipid peroxidation by measuring the quantities of malondialdehyde (MDA) generated during cellular lipid peroxidation. To summarize, the cell lysates were centrifuged at 10,000 g for 10 minutes at 4°C. The supernatants were then treated with an equal volume of TBA solution (0.375% TBA, 15% trichloroacetic acid, and 0.25 N HCl) and heated for 15 minutes in a boiling water bath before centrifugation at 10,000 g for 5 minutes. Finally, the supernatant’s absorbance was measured at 535 nm. Fold change over control is used to express the change in MDA level [69].

### Determination of mitochondrial membrane potential

Mitochondrial membrane potential (ΔΨm) was measured using a cationic dye, JC-1. A 6-well plate was seeded with 4 ×10^5^ HeLa cells per well. After 24 h, cells were treated with different concentrations of DCF and incubated for 24 h. Then cells were washed with PBS (pH 7.4) and stained with 20 μM of JC-1 for 30 min at 37 °C. PBS (pH 7.4) was used to wash the cells, and the cell suspension was analyzed by FACS Verse (Becton-Dickinson, USA).

### Determination of apoptosis by Annexin V/PI staining

The Annexin V/PI dual staining assay was used to detect apoptotic cells. Initially, 4 x10^5^ cells were plated into each well of a 6-well plate and cultured for 24 h. Then cells were treated with different concentrations of DCF, incubated for 24 h, and washed twice with PBS (pH 7.4) at room temperature. Cells were resuspended in 100 µL annexin-V binding buffer. Subsequently, 5 µL annexin V-FITC and 5 µL propidium iodide were added, and the cells were incubated at room temperature for 15 minutes in the dark. FACS analysis was performed, and the fold change of apoptotic and necrotic populations was estimated.

### Caspase 3 activity assay

4 x 10^5^ HeLa cells were seeded in each well of a 6-well plate. After 24 h, cells were treated with different concentrations of DCF and incubated for 24 h. Caspase 3 activity was measured by a caspase-3 activity assay kit (Elabscience) following the manufacturer’s instructions.

### Determination of the docking score for binding of DCF with enzymes in the glycolytic pathway

The three-dimensional structures of the glycolytic enzymes were obtained from RCSB-PDB, and the 3D structure of DCF was obtained from PubChem. The docking score of the individual enzyme drug complex was determined using LeadIT 2.3.2 software. All the grid values, reference ligands, and other parameters were kept constant during docking studies. Five docking scores were determined for each DCF-enzyme complex.

### Prediction of the binding sites on the LDH-A/DCF interaction

Molecular docking using DCF, pyruvate, and sodium oxamate (inhibitor of LDH-A) with lactate dehydrogenase-A (LDH-A) was performed by the AutoDock Vina software. The crystal structure of LDH-A (PDB ID: 1i10) was collected from the RCSB PDB. During docking, all the parameters were kept constant.

### Binding score determination of LDH-A/DCF and LDH-B/DCF complex

The *in-silico* binding score of the LDH-A/DCF and LDH-B/DCF complexes was determined using the AutoDock software to check the preliminary binding affinity of DCF towards LDH-A and LDH-B. The 3D structures of LDH-A and LDH-B were obtained from the RCSB PDB. PDB ID of LDH-A is 1i10 and LDH-B is 1T2F. During docking, all the parameters were kept constant.

### Measurement of intracellular LDH-A activity

HeLa, NKE and HaCaT cells were plated in 6-well plates at a density of 0.3 x 10^6^ cells per well. After reaching 70% confluency, treatment was done with two different DCF concentrations (for HeLa cells). 100 μM sodium oxamate was used as a positive control. After 24 h of incubation with DCF and sodium oxamate, cells were washed with PBS (pH 7.4). Samples were sonicated and centrifuged at 8000 rpm for 15 minutes. The supernatant was collected, and protein estimation was done using Bradford’s reagent (SRL). 50 mM Tris buffer (pH 7.6), 1.8 mM sodium pyruvate, 150 μM NADH, and cell lysates with equal protein content (50 μg) were added. After proper mixing, the change in OD was measured after 3 minutes of reaction. LDH activity was directly proportional to ΔOD. Where ΔOD = change in OD over 3 min.

### Protein extraction and western blot analysis

HeLa cells were lysed using RIPA lysis buffer (50 mM Tris-HCl pH 7.4, 150 mM NaCl, 0.1% SDS, 1% NP40, 0.5% sodium deoxycholate, 2 mM EDTA, and protease inhibitor cocktail), followed by a 30 min incubation on ice. The cell lysate was centrifuged at 13000g for 15 min at 4 °C. The pellet was discarded, and the supernatant was used for protein estimation using Bradford reagent. Equal amounts of protein were loaded onto an SDS-PAGE denaturing gel and transferred to a PVDF membrane. After successful transfer, 5% non-fat milk was used as a blocking reagent, and the membrane was blocked for 1 h at room temperature, followed by incubation with the primary (1:1000 dilution) and secondary antibodies (1:5000 dilution). Visualization of the expression of the target protein was performed using SuperSignal™ West Pico PLUS Chemiluminescent Substrate (Thermo Fisher), followed by exposure of the blot onto an X-ray film. Densitometric studies were performed using ImageJ software, and the expression of the target protein was normalized over the control.

### Measurement of pure LDH-A activity

The enzymatic activity of pure LDH-A was estimated by monitoring the oxidation of NADH at 340 nm for 3 minutes. The final reaction mixture contained 50 mM tris buffer (pH 7.6), 1.8 mM sodium pyruvate, 150μM NADH, and 0.015 U.ml^-1^ pure muscle LDH-A (Sigma Aldrich). Two different DCF concentrations were used. Sodium oxamate was used as a positive control. LDH activity was directly proportional to ΔOD. Where ΔOD = change in OD over 3 min.

### Determination of pyruvate concentration (*in-vitro*)

The *in-vitro* pyruvate concentration of the sample was measured using the modified Schwimmer and Weston method [70]. At first, LDH-A activity in both treated and untreated samples was measured with different concentrations of DCF and oxamate (the positive control), as described earlier. After 3 min of the reaction, 100 μl DNPH (100 μM) was added to each sample. Then 100 μl of 1M HCl was added to each sample and incubated at 37 °C in a water bath for 10 min. Then 100 μl of freshly prepared 1.5 M NaOH was added, and the absorbance was measured at 515 nm using a spectrophotometer. Different known concentrations of sodium pyruvate were used to make a standard curve for the estimation of the pyruvate concentration in the unknown samples.

### Determination of intracellular LDH-B activity

HeLa cells were plated in 6-well plates at a density of 0.3 x 10^6^ cells per well. After reaching 70% confluency, treatment was done with different concentrations of DCF. After 24 h of incubation with DCF, cells were washed with PBS. Samples were sonicated and centrifuged at 8000 rpm for 15 minutes. The supernatant was collected, and protein estimation was done using Bradford’s reagent. As an enzyme source, 50 μg of total protein from cell lysates was employed for intracellular LDH-B activity experiments. LDH-B activity was measured in cell lysates using the LDH activity assay kit (Elabscience) according to the manufacturer’s instructions.

### Determination of the reaction kinetics of LDH-A

The reaction kinetics of LDH-A were estimated by determining the change in the absorbance of NADH at 340 nm over 3-minute intervals. The enzyme kinetics of LDH-A were determined in two separate experiments. In the first experiment, NADH concentration was kept constant (150μM) and the reaction mixture contained 50 mM tris buffer (pH 7.6),175 μM DCF, 0.015 U.ml^-1^ pure LDH-A, and a range of sodium pyruvate (1–5 mM). In the second experiment, all the reagent concentrations were the same except the sodium pyruvate concentration, which was kept constant at 1 mM, and a concentration range of NADH (0.5 μM-75 μM) was used.

### Determination of the fluorescence intensity of LDH-A

100 μg of pure LDH-A was taken in 50 mM phosphate buffer at pH 7.4, and different concentrations of DCF were added at 25 °C and incubated for 25–30 min. The fluorescence intensity was measured using a spectrofluorometer (Horiba) (the excitation wavelength was kept at 295 nm). Fluorescence emission was observed at 327 nm, and the quenching constant (K_q_) and binding constant (K_a_) of the complex were determined from the fluorescence quenching data.

### Circular dichroism (CD) spectroscopy

A CD spectrophotometer (J-815 JASCO, Bose Institute, India) with a 300 μl sample quartz cup was used to acquire the CD spectra of pure LDH-A in the far-UV region from 200 nm to 260 nm. Before obtaining the CD spectra, the LDH-A concentration was set to 1 mg/ml. LDH-A was incubated with 25 μM DCF for 10 minutes. The data was collected at 25 °C. Each individual sample was scanned three times.

### Determination of glucose uptake rate, pyruvate concentration, lactate production, NAD^+^ concentration, and ATP concentration

HeLa cells were seeded at a density of 1 x 10^4^ cells per well in a 96-well plate. Then cells were treated with different doses of DCF for 24 and 48 h. Here, sodium oxamate was used as a positive control. Glucose uptake rate, pyruvate concentration, extracellular lactate concentration, NAD^+^ concentration, and ATP concentration were determined using a glucose uptake assay kit (Elabscience), pyruvate assay kit (Elabscience), lactate assay kit (Elabscience), ATP assay kit (Elabscience), and NAD^+^ concentration assay kit (Cayman) as per the manufacturer’s instructions.

### Induction of CoCl_2_-induced hypoxia in HeLa cells

HeLa cells were cultured in a 6-well plate at a density of 0.3 x 10^6^ cells per well. After 24 h of incubation, cells were treated with different concentrations of CoCl_2_ and incubated for 24 h at 37 °C in a 5% CO_2_ atmosphere.

### Small interfering RNA (siRNA) transfection

Lipofectamine RNAiMAX transfection reagent was used to transfect predesigned small interfering RNA (siRNA) against LDH-A (5’GGAGAAAGCCGUCUUAAUU3’,5’ GGCAAAGACUAUAAUGUAA 3’), AMPK α1 (5’ GCAUAUGCUGCAGGUAGAU 3’) and control scrambled siRNA (sc-37007) in HeLa cells by using the manufacturer’s protocol. All transfection experiments were carried out after 48 h of incubation with siRNA.

### Statistical analysis

All data are presented as mean ± SD and are based on at least three independent experiments. A P value of 0.05 was regarded as significant. The Student’s t test was employed to compare group differences.

## Abbreviation

LDH-A: Lactate dehydrogenase -A
DCF: Diclofenac sodium
NSAID: Non-steroidal anti-inflammatory drug
AMPK: AMP activated protein kinase
AMP: Adenosine mono-phosphate
ADP: Adenosine di-phosphate
NADH: Nicotinamide adenine dinucleotide (Reduced)
NAD^+^: Nicotinamide adenine dinucleotide (oxidised)
HIF-α: Hypoxia induced factor-α
ATP: Adenosine triphosphate
COX-2: Cyclooxygenase-2
NKE: Normal kidney epithelial
PI: Propidium iodide
DCFDA: 2’-7’-Dichlorodihydrofluorescein diacetate
FACS: Fluorescence-activated cell sorting
JC-1: 5,5,6,6’-tetrachloro-1,1’,3,3’ tetraethylbenzimi-dazoylcarbocyanine iodide
PDB: Protein data bank
OD: Optical density
SD: Standard deviation
DMEM: Dulbecco’s Modified Eagle’s Medium
CD: Circular dichroism
UV: Ultra-violet
ROS: Reactive oxygen species
DNA: Deoxyribonucleic acid
PUFA: Polyunsaturated fatty acids
MDA: Malondialdehyde
OXPHOS: Oxidative phosphorylation
BSA: Bovine serum albumin
TBARS: Thiobarbituric acid reactive substance
AICAR: 5-Aminoimidazole-4-carboxamide ribonucleoside
TBA: Thiobarbituric acid
PBS: Phosphate buffered saline
SDS: Sodium Dodecyl Sulfate
SV: Stern Volmer
EDTA: Ethylenediamine tetra acetic acid
PAGE: Polyacrylamide gel electrophoresis
DNPH: 2,4-dinitrophenylhydrazine

## Data availability statement

All raw data and materials will be made available following a reasonable request.

## Author contributions

Conceptualization of the project was done by SG and RS and AM. Acquisition of data was done by AM. Data analysis and interpretation was done by SG, RS and AM. AM prepared the original draft which was edited by SG and RS. SG and RS are corresponding authors of this manuscript. All approve the final version of the submitted manuscript.

## Funding and additional information

The fellowship of AM has been funded by Council of Scientific & Industrial Research [CSIR File no-09/028(1112)/2019-EMR-I]. This study was supported by a grant from the Science and Engineering Research Board (SERB), departmental research grant of the University of Calcutta and DST-FIST and DST-TARE (Sanction # TAR_2018_000336)

## Acknowledgements

We thank the Council of Scientific & Industrial Research (CSIR), Science and Engineering Research Board (SERB), Department of Biophysics Molecular biology and Bioinformatics University of Calcutta, west Bengal India; Department of Biotechnology Haldia Institute of technology, West Bengal, India. The fellowship of the author has been funded by Council of Scientific & Industrial Research (CSIR). We want to acknowledge Dr. Soumalee Basu, department of microbiology, University of Calcutta for accessing her lab for docking experiments and Central Instrument Facility (CIF), Bose Institute Kolkata for conducting FACS and CD experiments.

