## supporting information for "Inhibition of Lactate Dehydrogenase A (LDH-A) by Diclofenac Sodium Induces Apoptosis in HeLa Cells by Activation of AMPK"

**Supplementary Figures**

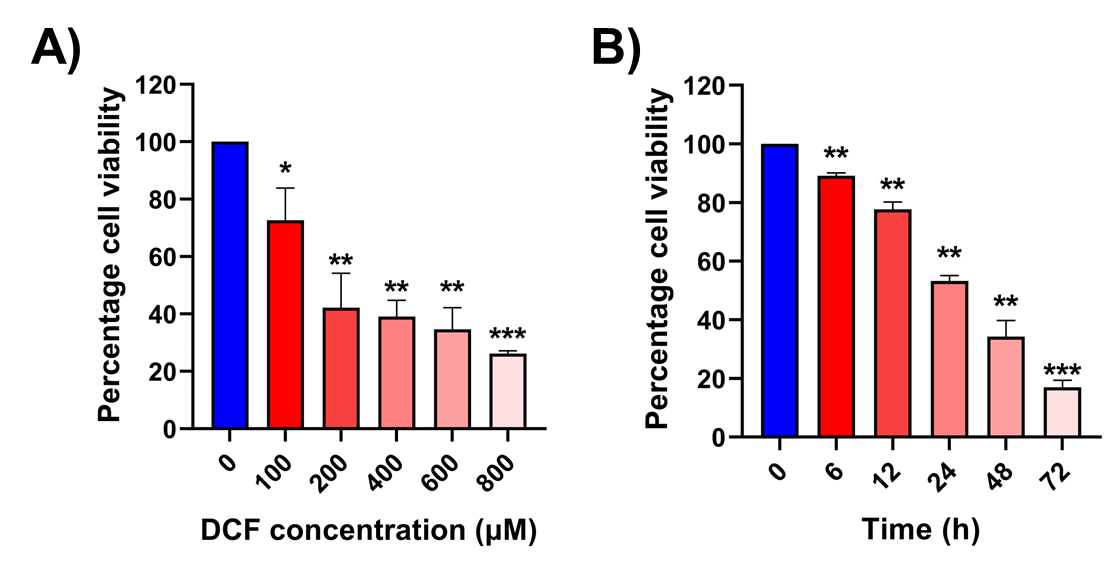

**Supplementary figure S1. Effect of DCF on the cell viability of HCT-116 cells.** A. The percentage of cell viability was estimated with different concentrations of DCF (0–800 μM) in HCT-116 cells for 24 h using the trypan blue exclusion assay. B. Cell viability was estimated with 150 μM DCF for different time points (0–72 h) in HCT-116 cells. ∗P<0.05, ∗∗P<0.01, ∗∗∗P<0.001 compared to the control group. All data represent mean ±SD of at least three independent experiments.

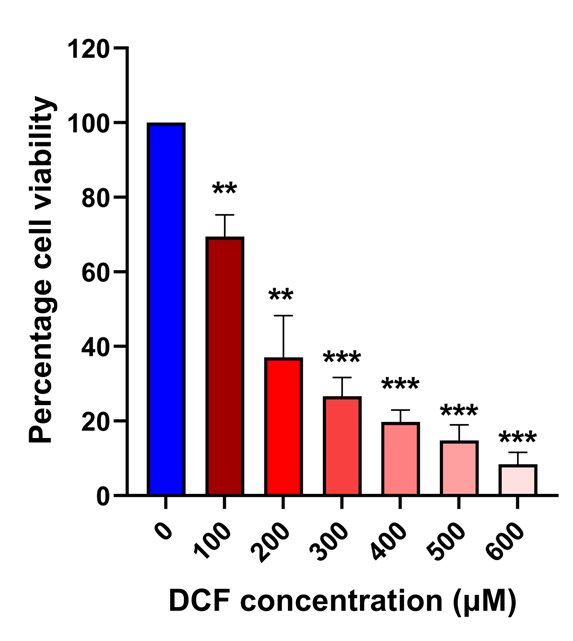

**Supplementary figure S2. Effect of DCF on the cell viability of MCF-7 cells.** The percentage of cell viability was estimated with different concentrations of DCF (0–600 μM) in MCF-7 cells for 24 h using the trypan blue exclusion assay. ∗P<0.05, ∗∗P<0.01, ∗∗∗P<0.001 compared to the control group. All data represent the mean ±SD of at least three independent experiments.

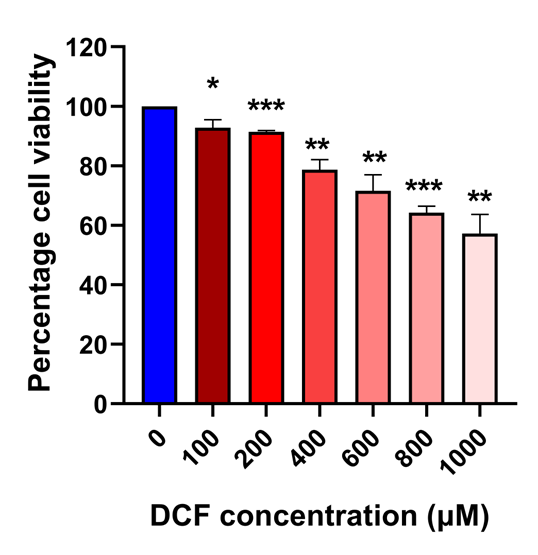

**Supplementary figure S3. Effect of DCF on the cell viability of HaCaT cells.** The percentage of cell viability was estimated with different concentrations of DCF (0–1000 μM) in HaCaT cells for 24 h using the trypan blue exclusion assay. ∗P<0.05, ∗∗P<0.01, ∗∗∗P<0.001 compared to the control group. All data represent the mean ±SD of at least three independent experiments.

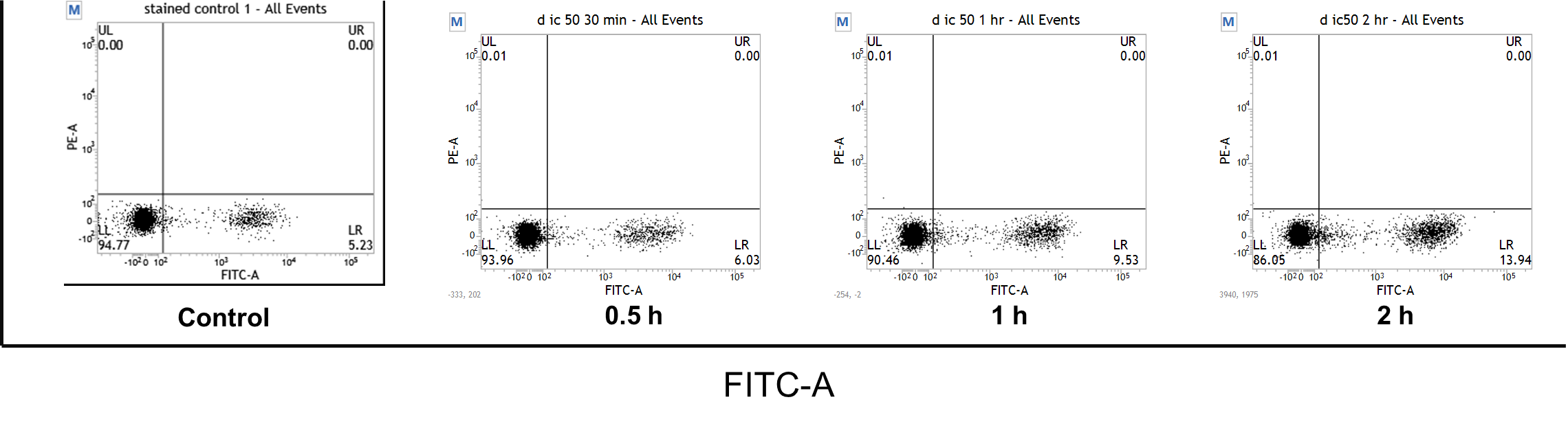

**Supplementary figure S4. Percentage distribution of H2DCFDA-stained HeLa cells.** HeLa cells were treated with 175 μM of DCF and incubated for different time points. A. untreated control; DCF treatment for B. 0.5 h, C. 1 h, and D. 2 h. ROS levels were measured by H2DCFDA staining followed by FACS analysis. The percentage distribution of cells in the LR (lower right) quadrant indicates the population of DCFDA-positive cells after incubation with DCF at the indicated time points.

**A)**

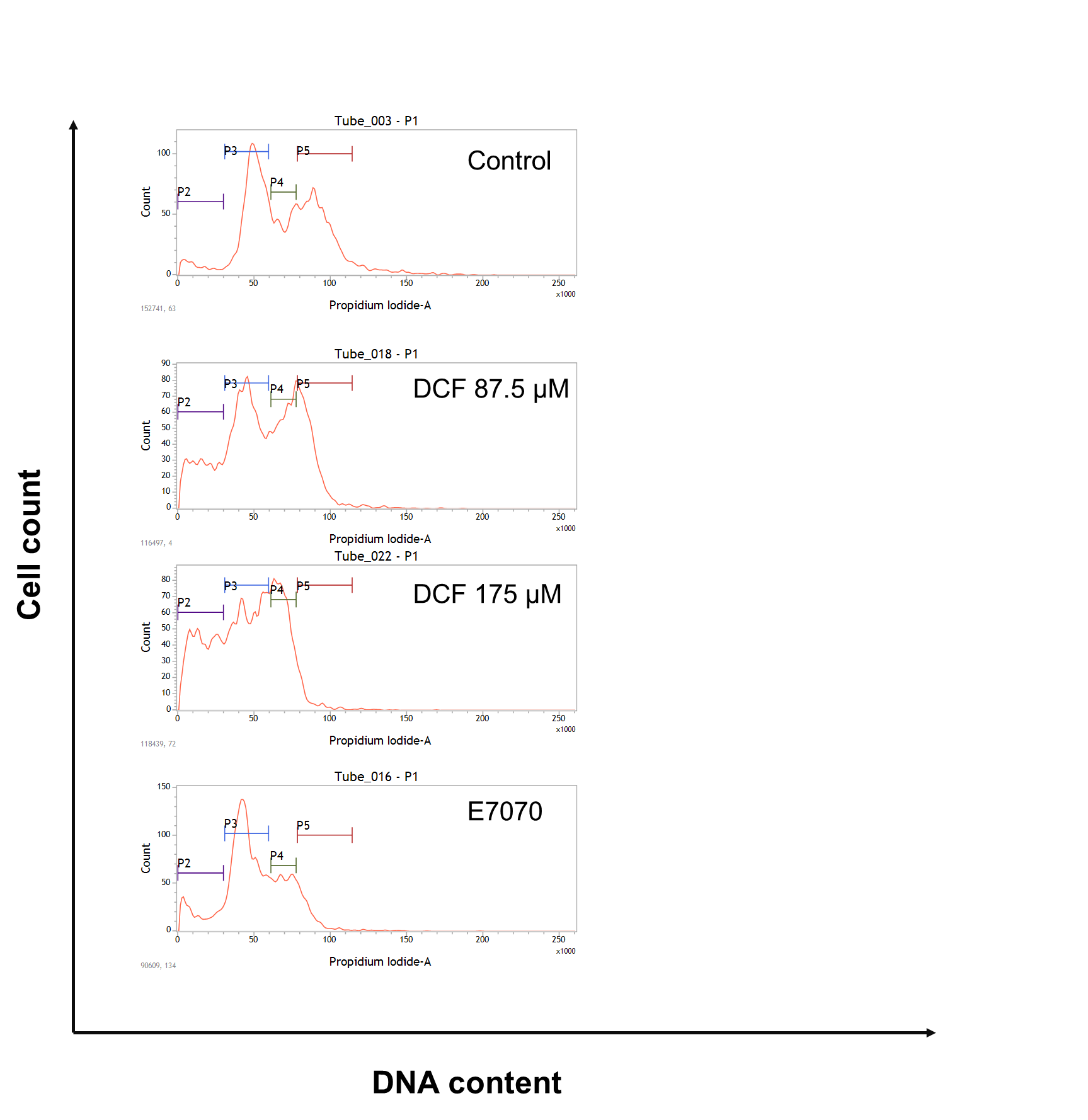

**B)**

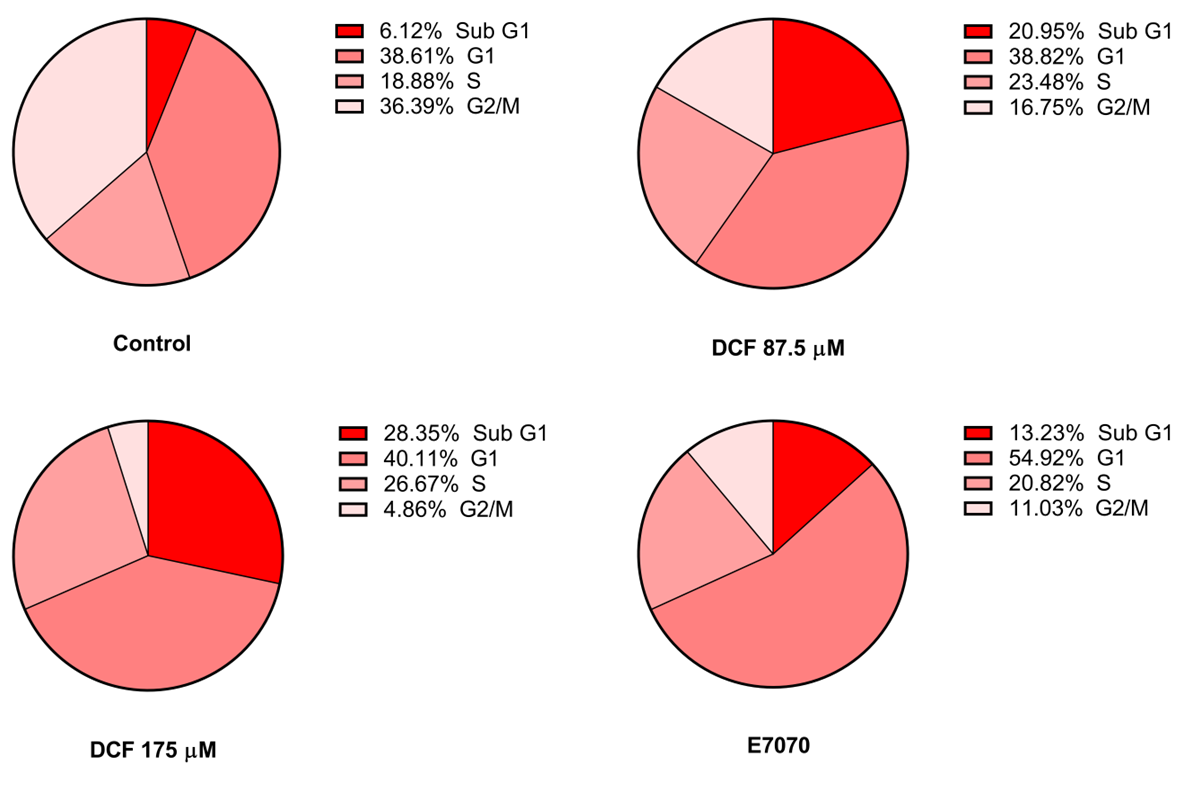

**Supplementary figure S5. Effect of DCF on cell cycle progression of HeLa cells.** A. Effect of different concentrations of DCF and E7070 (positive control) on the cell cycle distribution using PI staining followed by FACS analysis B. The pie chart represents the percentage of cell population in Sub-G1, G1, S, and G2/M phases obtained from Fig. S5A.

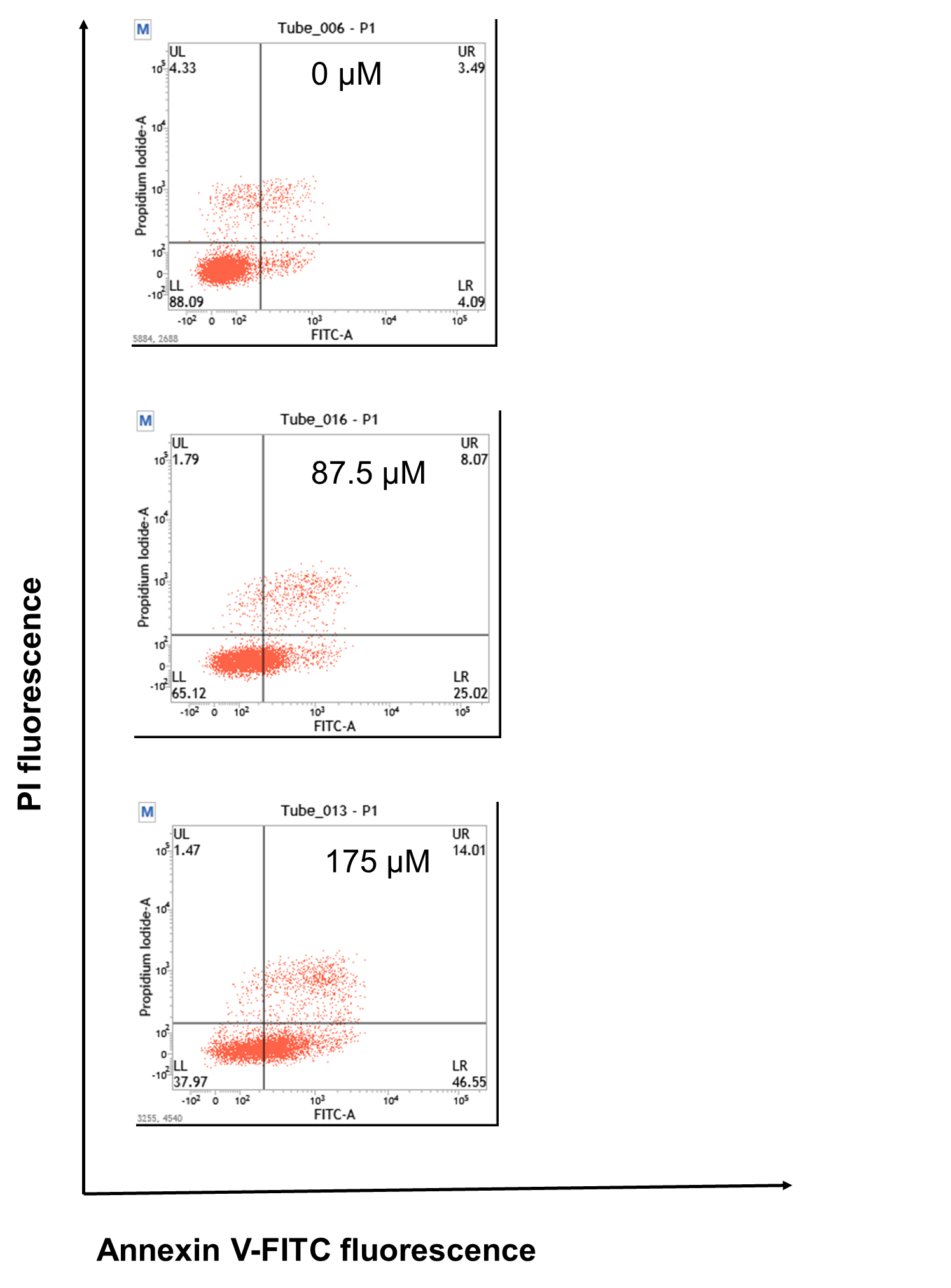

**Supplementary figure S6. Effect of DCF on different phases of apoptosis.** HeLa cells were treated with DCF (indicated concentrations) for 24 h. Apoptotic (annexin-V^+^/PI^-^ for early apoptotic and annexin-V^+^/PI^+^ for late apoptotic) and necrotic (annexin^-^V-/PI^+^) cells were analyzed by flow cytometry (details are described in the methods section). Each quadrant indicates the percentage population of cells in different stages of apoptosis (LR and UR quadrants) and necrosis (UL quadrant).

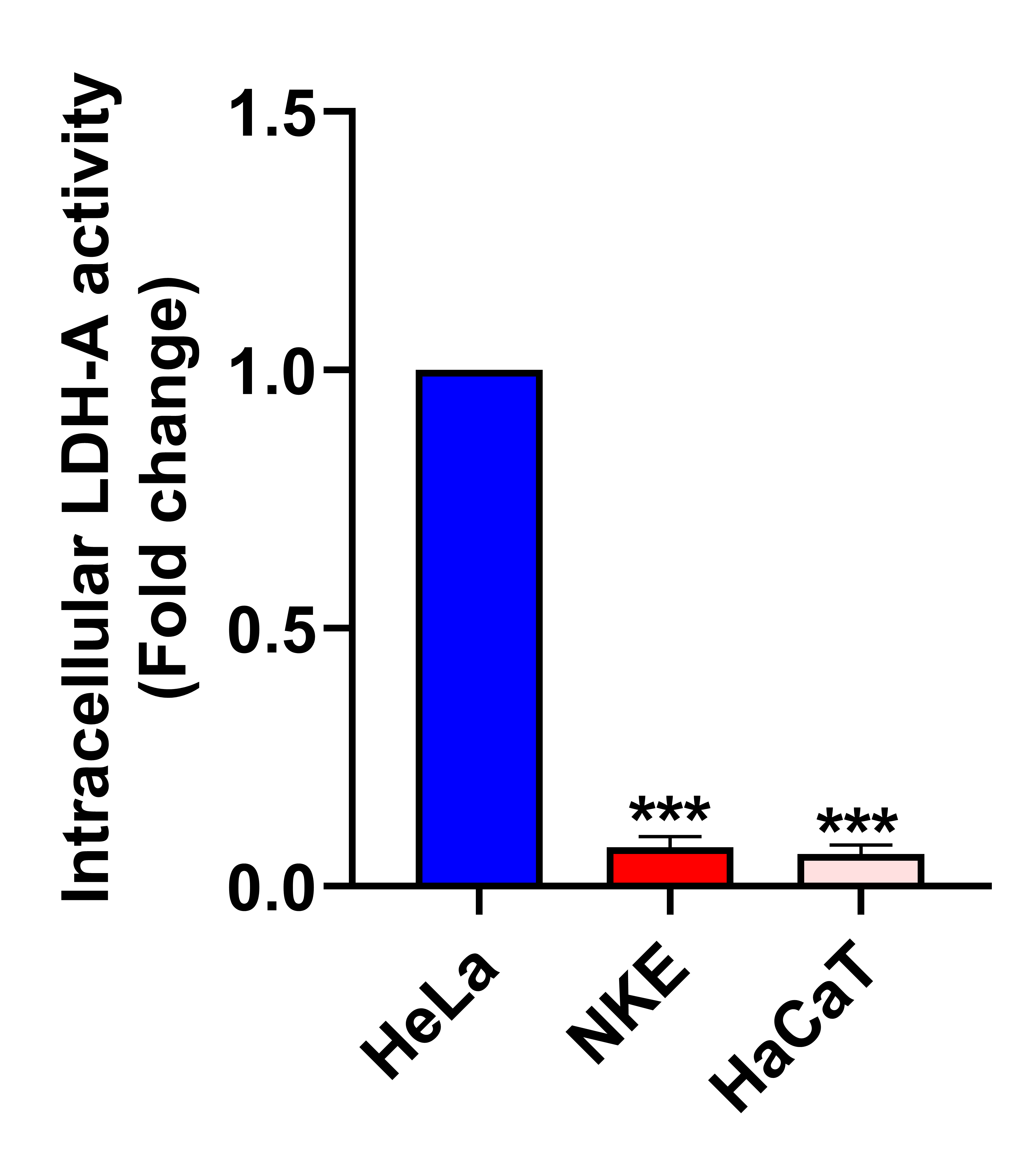

**Supplementary figure S7. Intracellular LDH-A activity in HeLa, NKE, and HaCaT cells.** Intracellular LDH-A activity in cells was estimated by measuring the change in OD using a spectrophotometer. The bar diagram indicates the fold change in the LDH-A activity of NKE and HaCaT cells compared to HeLa cells. All the bar graphs represent the mean ± SD of three independent experiments where ∗P<0.05, ∗∗P<0.01, and ∗∗∗P<0.001 when compared to the LDH-A activity in HeLa cells.

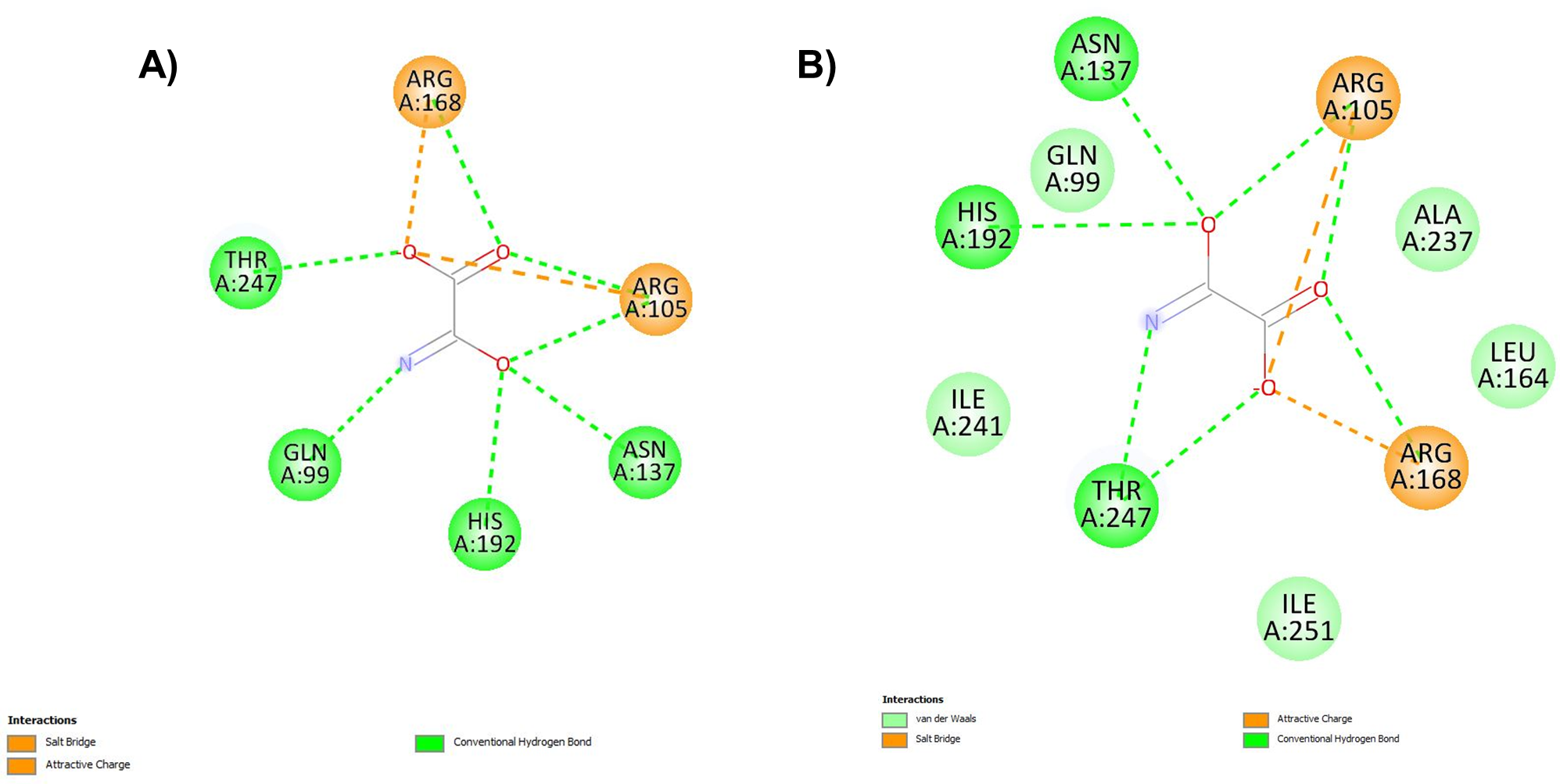

**Supplementary figure S8. Validation of AutoDock using oxamate and LDH-A.** AutoDock was validated by redocking oxamate with the crystal structure of LDH-A (PDB ID (1i10) of LDH-A, which was bound with oxamate. A. Reported interactions of oxamate with the crystal structure of LDH-A. B. Interactions of the re-docked oxamate with LDH-A using the AutoDock software. Dotted lines indicate the different types of interactions of oxamate with specific amino acids in LDH-A.

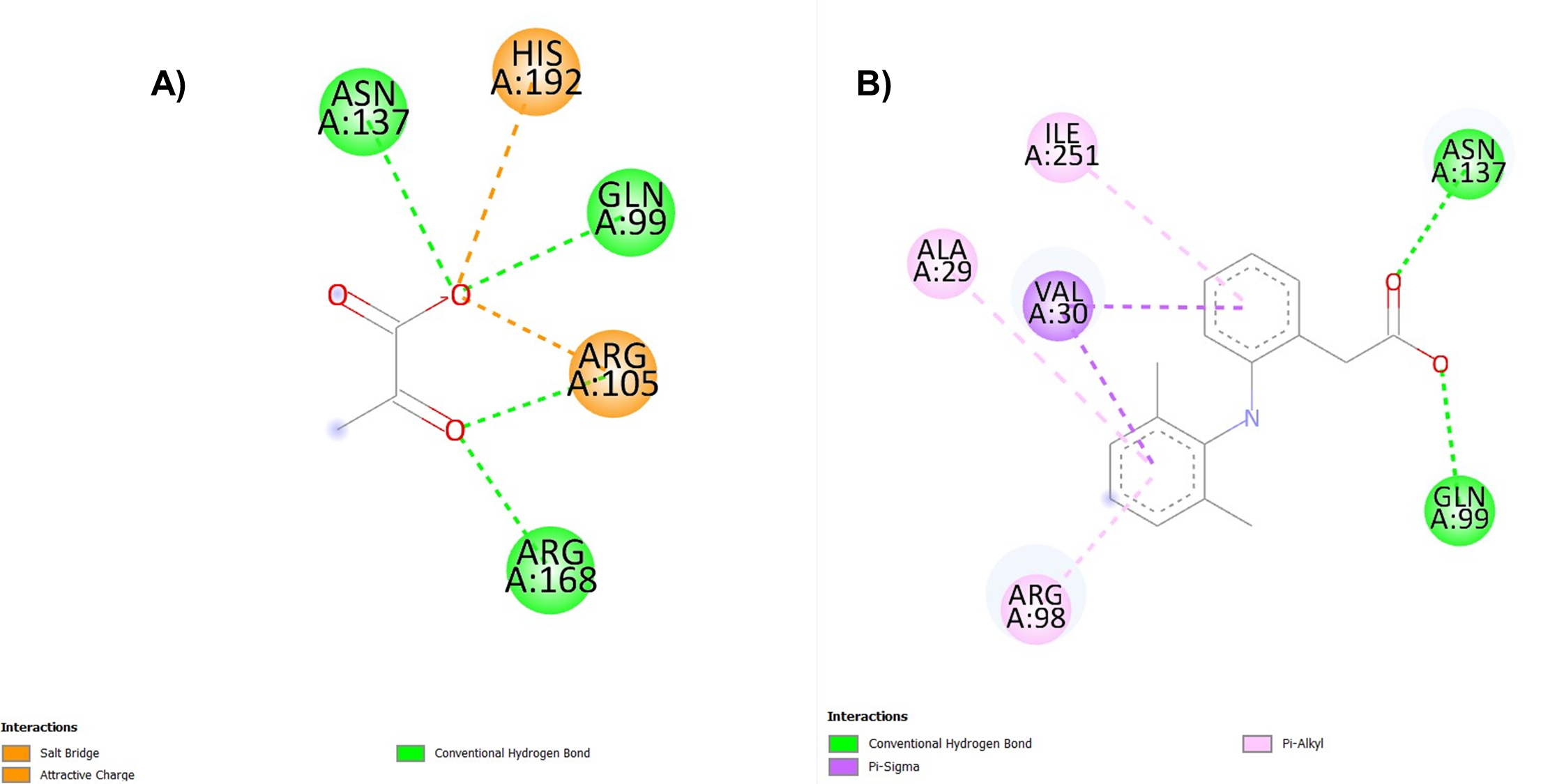

**Supplementary figure S9. DCF interacts with LDH-A in the vicinity of the substrate (pyruvate) binding site.** A. Residue-level interactions of pyruvate (a substrate of LDH-A) with LDH-A using AutoDock B. Residue-level interactions of DCF with LDH-A. Each dotted line indicates the different interactions of specific ligands with LDH-A. Each type of interaction is listed in the diagram.

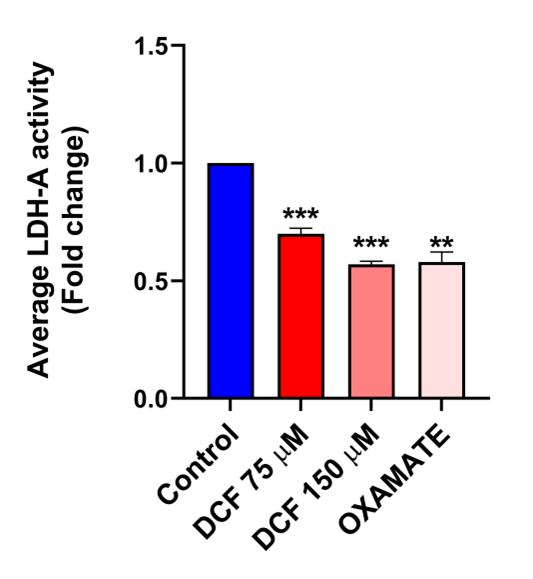

**Supplementary figure S10. DCF inhibits intracellular LDH-A activity in HCT-116 cells.** Intracellular LDH-A activity was measured with different concentrations of DCF in HCT-116 cells for 24 h. Oxamate was used as a positive control. The bar diagram indicates the fold change in the enzymatic activity over control. All the bar graphs represent the mean ± SD of three independent experiments where ∗P<0.05, ∗∗P<0.01, and ∗∗∗P<0.001 when compared to the control group.

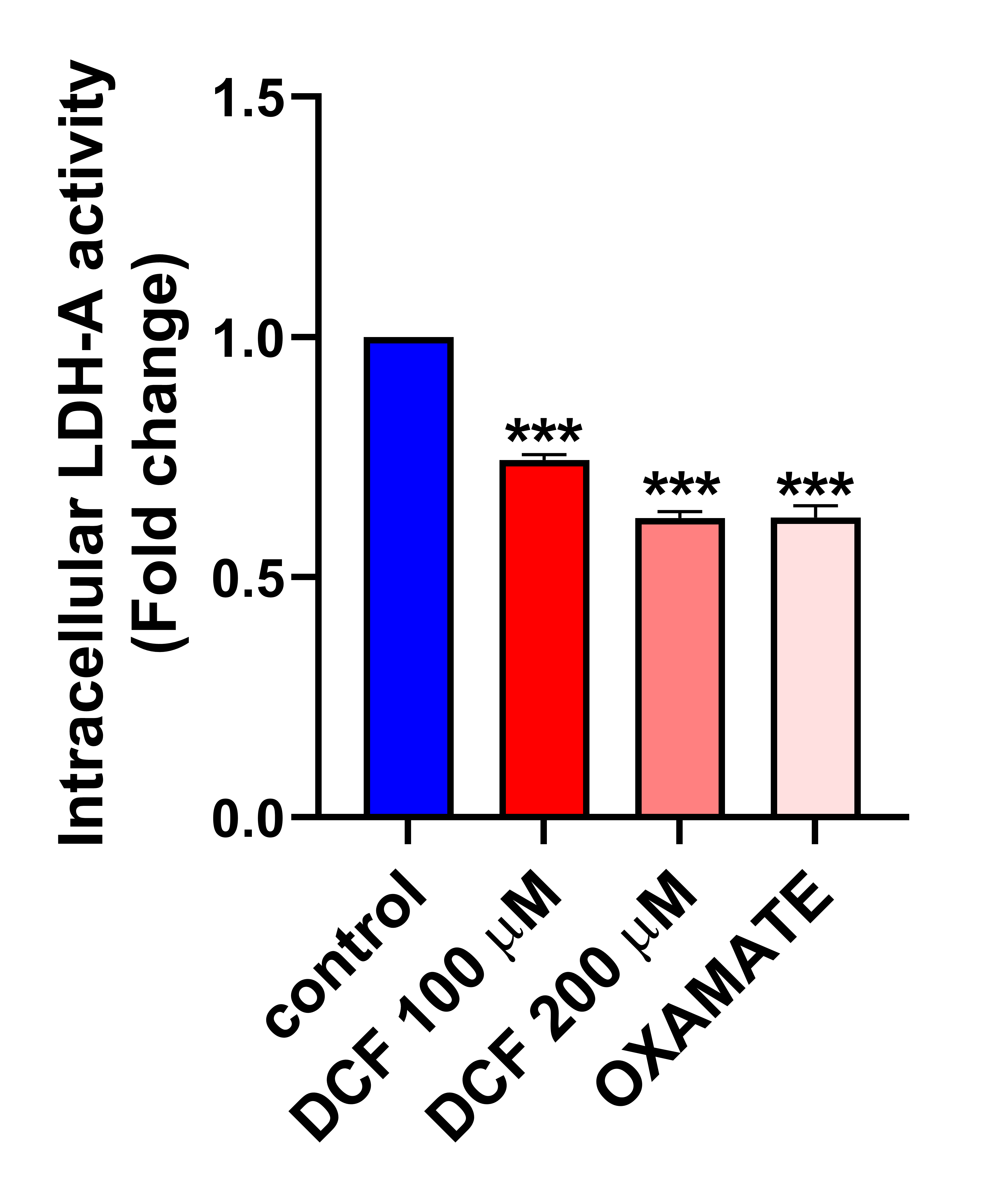

**Supplementary figure S11. DCF inhibits intracellular LDH-A activity in MCF-7 cells.** Intracellular LDH-A activity was measured with different concentrations of DCF in MCF-7 cells for 24 h. Oxamate was used as a positive control. The bar diagram indicates the fold change in the enzymatic activity over control. All the bar graphs represent the mean ± SD of three independent experiments where ∗P<0.05, ∗∗P<0.01, and ∗∗∗P<0.001 when compared to the control group.

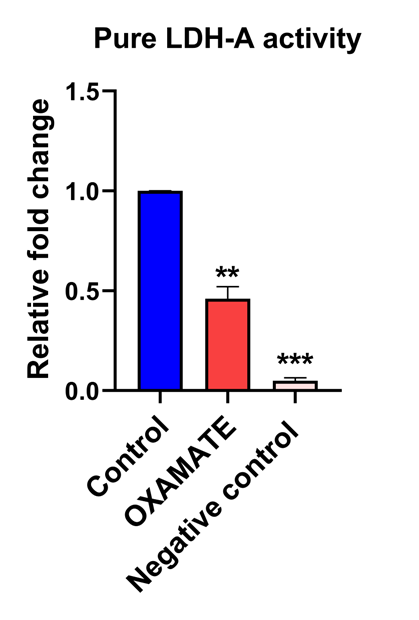

**Supplementary figure S12. Effect of oxamate and negative control on LDH-A activity in a cell-free system.**  LDH-A activity was measured in the presence of oxamate (a positive control) and in the absence of NADH (a negative control), where other parameters remained constant. All the bar graphs represent the mean ± SD of three independent experiments where ∗P<0.05, ∗∗P<0.01, and ∗∗∗P<0.001 when compared to the control group.

**Fig. S13**

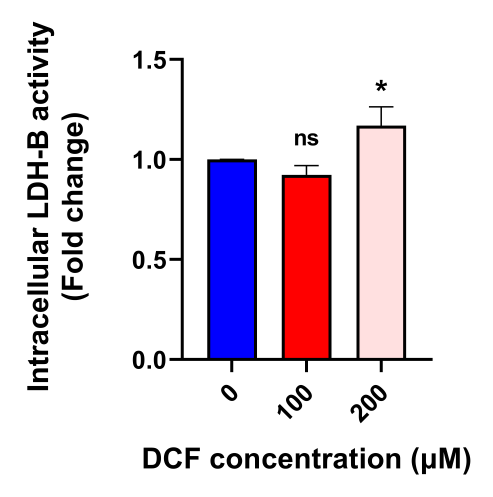

**Supplementary figure S13. DCF does not affect intracellular LDH-B activity in MCF-7 cells.** Intracellular LDH-B activity was measured in MCF-7 cells after treatment with different concentrations of DCF for 24 h. The bar diagram indicates the fold change in the enzymatic activity over control. All the bar graphs represent the mean ± SD of three independent experiments where ∗P<0.05, ∗∗P<0.01, and ∗∗∗P<0.001 0.001, ns: non-significant were compared to the control group.

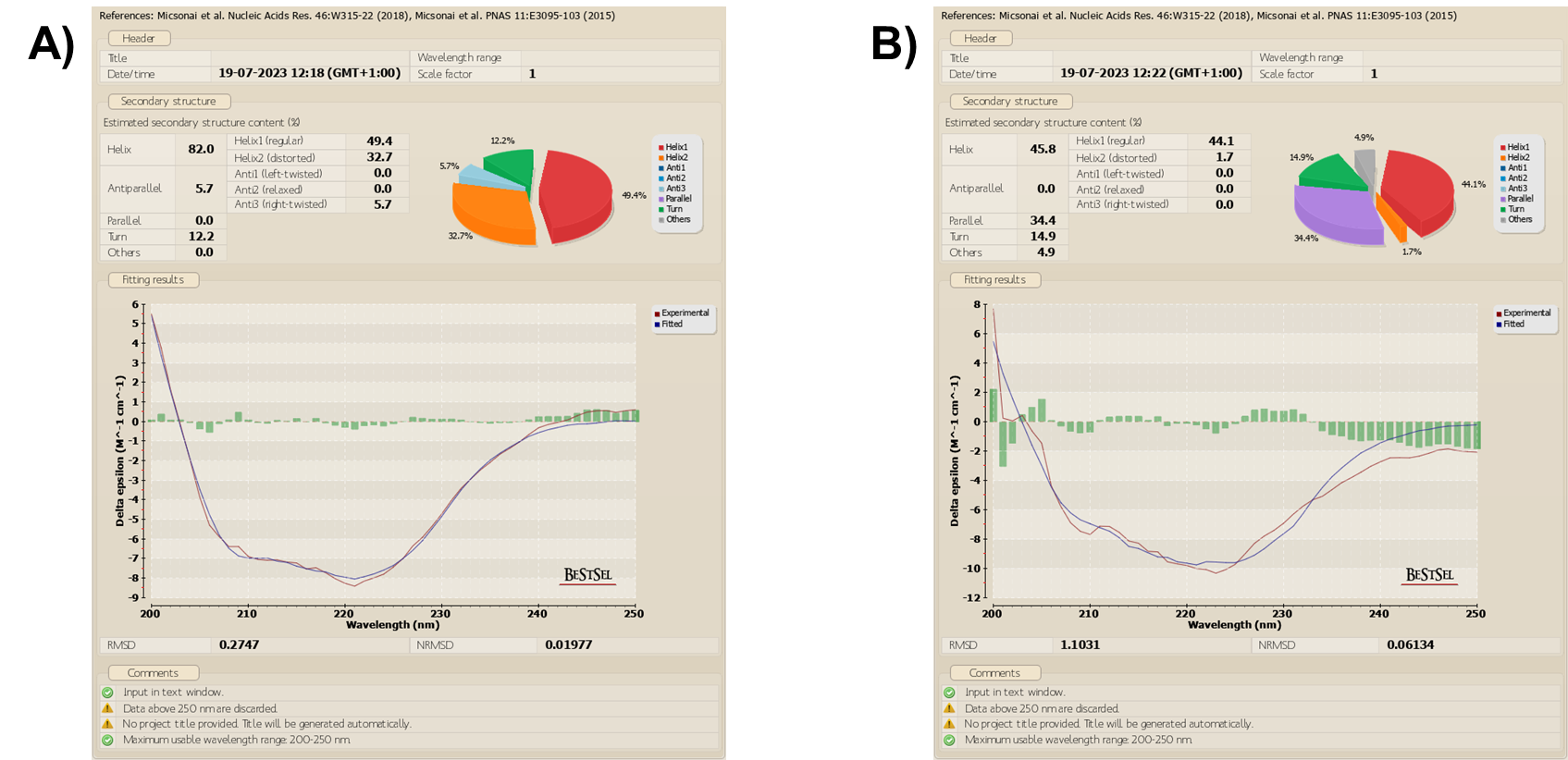

**Supplementary figure S14. Alternation of the secondary structure of LDH-A due to interaction with DCF.** Effect of DCF A. Control (0 μM DCF) and B. 25 μM on the secondary structures of LDH-A obtained from CD spectroscopy analysis. The percentage distribution of protein secondary structures was determined by fitting the CD spectroscopy results using protein circular dichroism spectra analysis software (BeStSel).

**Fig. S15**

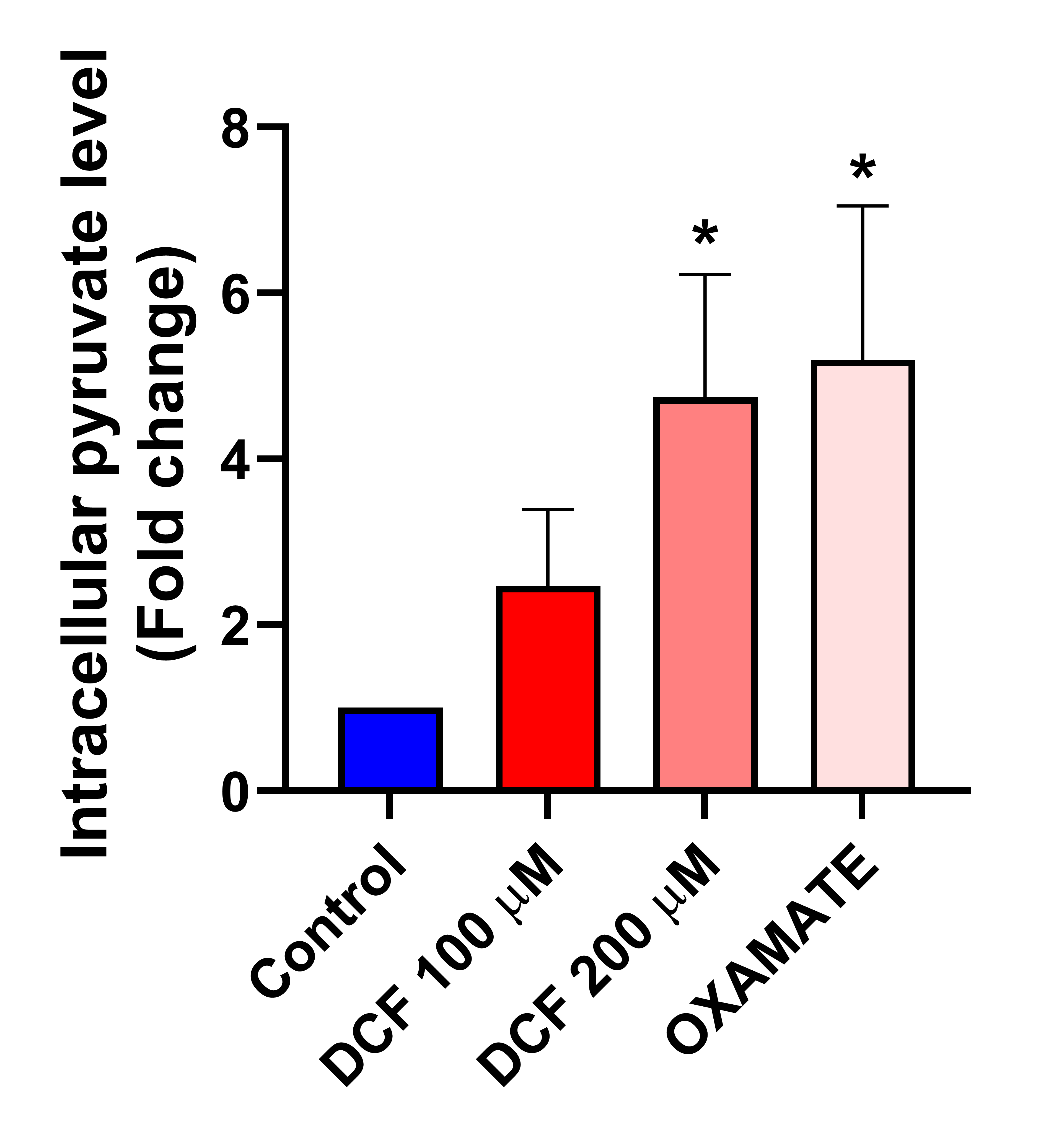

**Supplementary figure S15. DCF leads to the accumulation of intracellular pyruvate in MCF-7 cells.** MCF-7 cells were treated with different concentrations of DCF for 24 h, with oxamate as a positive control. The bar graph represents the mean ± SD of three independent experiments. ∗P<0.05, ∗∗P<0.01, and ∗∗∗P<0.001 compared to the control group

**Fig. S16**

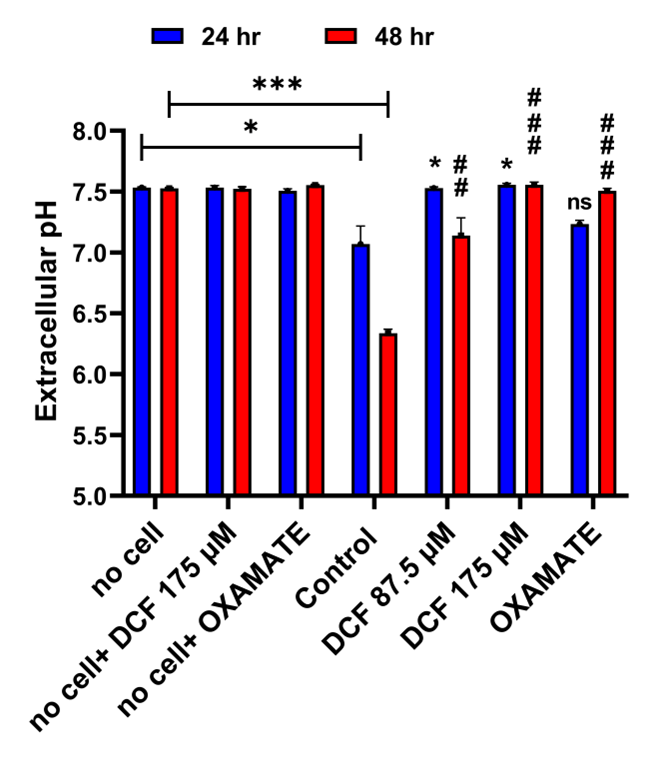

**Supplementary figure S16. DCF alters extracellular pH levels in HeLa cells.** HeLa cells were treated with different concentrations of DCF for 24 h, with oxamate as a positive control. The bar graph represents the mean ± SD of three independent experiments. No cell, no cell+ DCF, and no cell+ oxamate were used as negative controls. ∗P<0.05, ∗∗P<0.01, and ∗∗∗P<0.001 compared to the control group for 24 h. #P<0.05, ##P<0.01, and ###P<0.001 compared to the control group for 48 h.

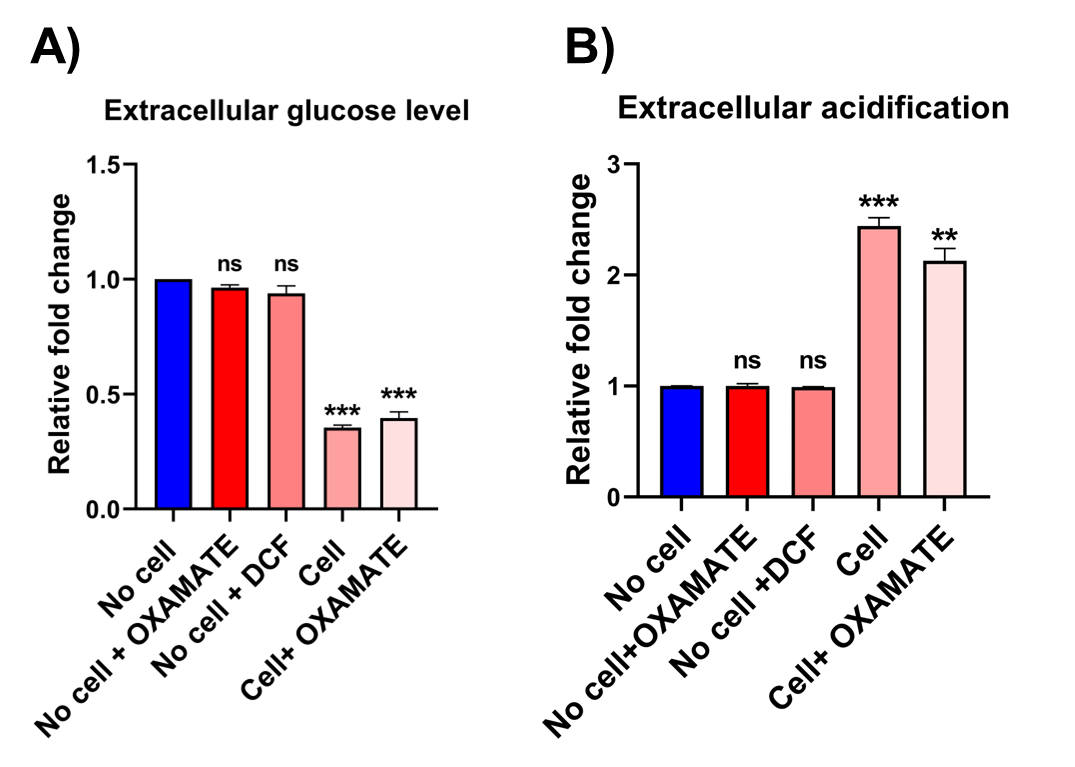

**Supplementary figure S17. Effect of positive and negative controls on extracellular glucose levels and extracellular acidification in HeLa cells.** A, Extracellular glucose level, and B, Extracellular acidification was measured in the presence of negative controls (no cell, no cell +oxamate, no cell + DCF) and in the presence of positive controls (cell, cell + oxamate). ∗P<0.05, ∗∗P<0.01, ∗∗∗P<0.001 and ns: non-significant compared to the control group

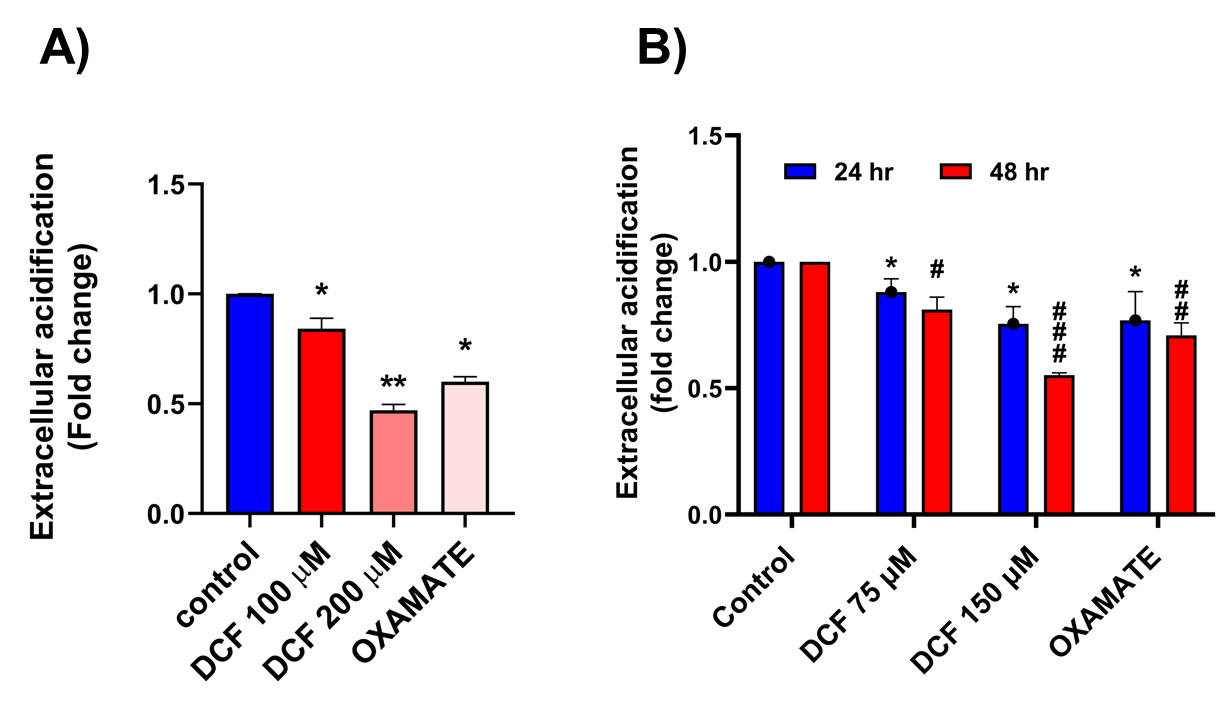

**Supplementary figure S18. DCF alters the extracellular acidification level in MCF-7 and HCT-116 cells.** Cells were treated with the indicated concentrations of DCF, and extracellular acidification was measured in A. MCF-7 cells at 24 h and in B. HCT-116 cells at 24 h and 48 h. Oxamate was used as a positive control. The bar graph represents the mean ± SD of three independent experiments. ∗P<0.05, ∗∗P<0.01, and ∗∗∗P<0.001 compared to the control group for 24 h. #P<0.05, ##P<0.01, and ###P<0.001 compared to the control group for 48 h.

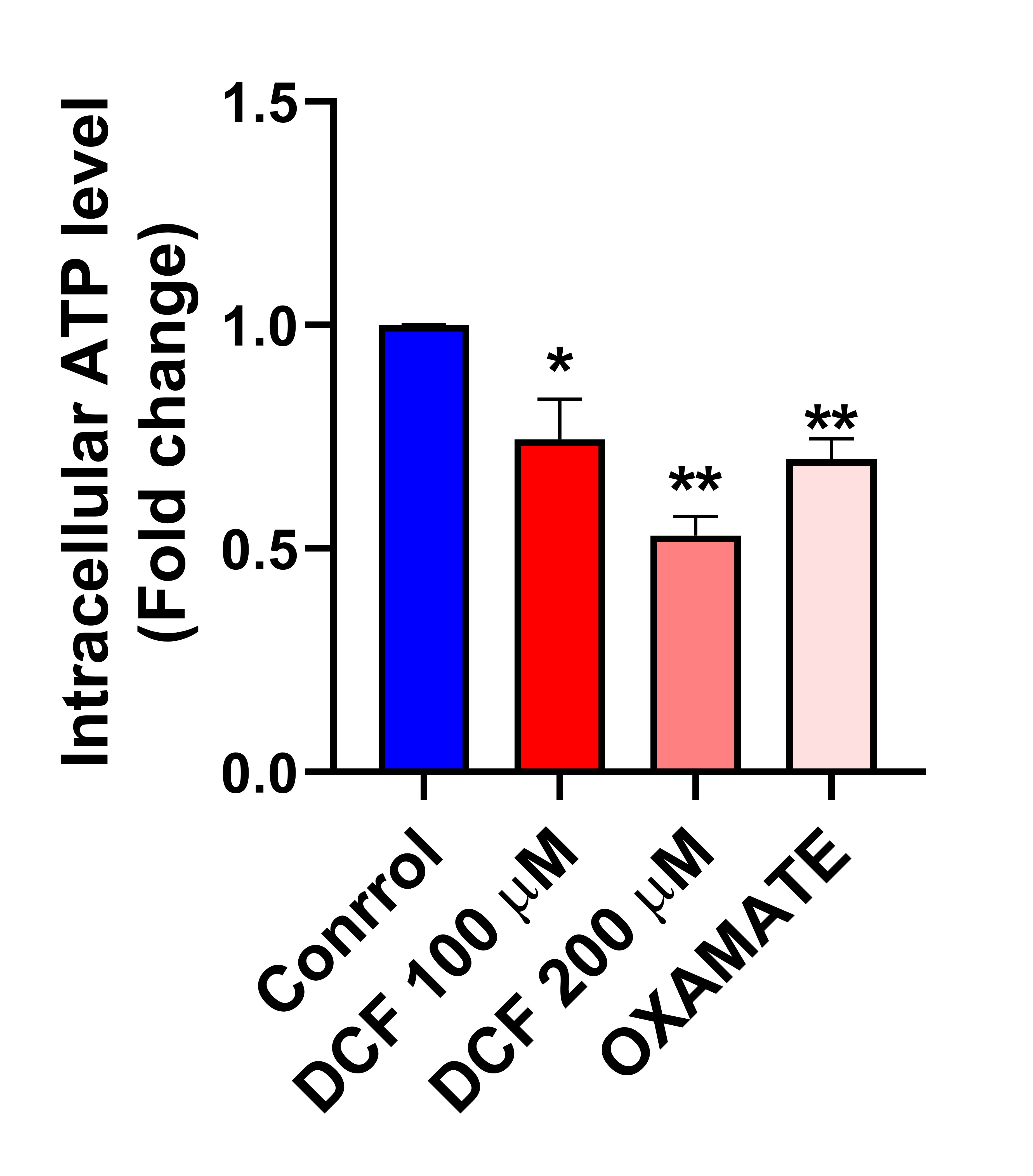

**Supplementary figure S19. DCF alters the intracellular ATP level in MCF-7 cells.** MCF-7 cells were treated with different concentrations of DCF for 24 h, and oxamate was used as a positive control. The bar graph represents the mean ± SD of three independent experiments. ∗P<0.05, ∗∗P<0.01, and ∗∗∗P<0.001 compared to the control group

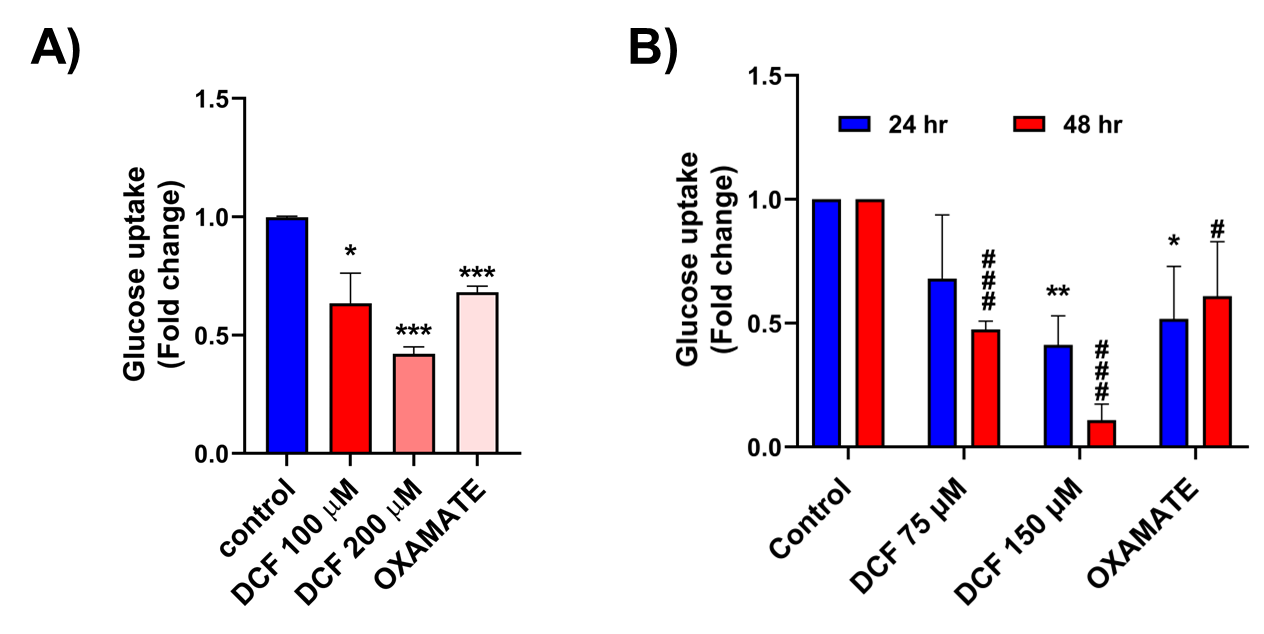

**Supplementary figure S20. DCF alters glucose uptake in MCF-7 and HCT-116 cells.** Effect of DCF on glucose uptake in A. MCF-7 cells at 24 h and B.  HCT-116 cells at 24 h and 48 h time points. Oxamate was used as a positive control. The bar graph represents the mean ± SD of three independent experiments. ∗P<0.05, ∗∗P<0.01, and ∗∗∗P<0.001 compared to the control group for 24 h. #P<0.05, ##P<0.01, and ###P<0.001 compared to the control group for 48 h.

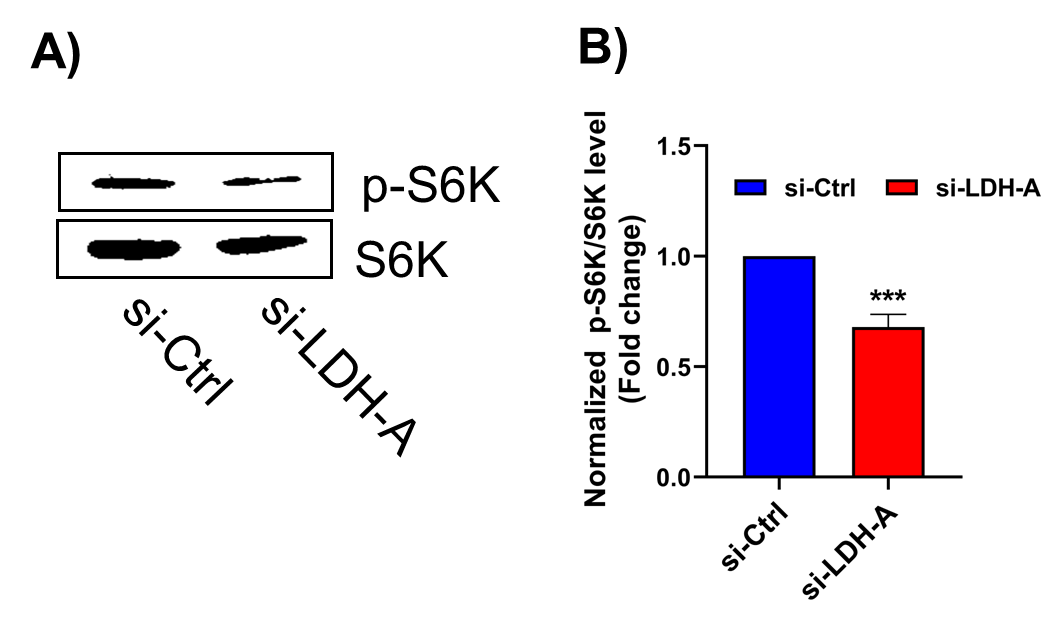

**Supplementary figure S21. Effect of si-LDH-A on the expression of p-S6K level in HeLa.** A. Western blot analysis of p-S6K and S6K expression in HeLa cells transfected with si-Ctrl and si-LDH-A for 48 h. B, Relative fold change of the normalized expression of p-S6K/S6K in si-Ctrl and si-LDH-A transfected cells at 48 h. ∗P<0.05, ∗∗P<0.01, and ∗∗∗P<0.001 compared to the control group.
